# CDKL5 modulates structural plasticity of excitatory synapses via liquid-liquid phase separation

**DOI:** 10.1101/2025.03.16.643581

**Authors:** Mingjie Li, Ziai Zhu, Dan Li, Jinchao Wang, Mingjie Zhang, Yongchuan Zhu, Jinwei Zhu, Zhi-Qi Xiong

**Affiliations:** Institute of Neuroscience, Center for Excellence in Brain Science and Intelligence Technology, Chinese Academy of Sciences, Shanghai 200031, China; University of Chinese Academy of Sciences, Beijing 100049, China; School of Future Technology, University of Chinese Academy of Sciences, Beijing 100049, China; School of Life Science and Technology, Shanghai Tech University, Shanghai 201210, China; Shanghai Center for Brain Science and Brain-Inspired Intelligence Technology, Shanghai 201210, China; Bio-X Institutes, Key Laboratory for the Genetics of Developmental and Neuropsychiatric Disorders, Ministry of Education, Shanghai Jiao Tong University, Shanghai 200240, China; Shanghai Key Laboratory of Psychotic Disorders, Shanghai Mental Health Center, Shanghai Jiao Tong University School of Medicine, Shanghai 200030, China; Department of Nephrology, Shanghai Sixth People’s Hospital Affiliated to Shanghai Jiao Tong University School of Medicine, Shanghai 200233, China; School of Life Sciences, Southern University of Science and Technology, Shenzhen 518055, China

**Author notes:** Correspondence to: Yongchuan Zhu; Jinwei Zhu; Zhi-Qi Xiong. The authors contributed equally to this work (Mingjie Li, Ziai Zhu).

## Abstract

Activity-dependent synaptic remodeling, essential for neural circuit plasticity, is orchestrated by central organizers within the postsynaptic density (PSD), including the scaffolding protein PSD95. However, the molecular mechanisms driving this process remain incompletely understood. Here, we identify CDKL5, a protein associated with a severe neurodevelopmental condition known as CDKL5 deficiency disorder (CDD), as a critical regulator of structural plasticity at excitatory synapses. We show that CDKL5 undergoes liquid-liquid phase separation (LLPS) in vitro and in cultured neurons, forming co-condensates with PSD95. This LLPS-driven process spatially organizes synaptic components, specifically enabling the synaptic recruitment of Kalirin7 to promote dendritic spine enlargement. Pathogenic mutations disrupt condensate formation by impairing the LLPS capacity of CDKL5, directly linking phase separation defects to the pathogenesis of CDD. Our findings reveal a crucial role for CDKL5 in synaptic plasticity and establish LLPS as a fundamental mechanism by which CDKL5 coordinates molecular events to reorganize PSD architecture during synaptic remodeling.

## Introduction

The cyclin-dependent kinase-like 5 (*CDKL5*) gene, located on the X chromosome, is genetically linked to CDKL5 deficiency disorder (CDD; OMIM 300203, 300672), a severe neurodevelopmental disorder characterized by intellectual disability, autistic-like features, global developmental delay, early-onset intractable epilepsy, hypotonia, and cortical visual impairment^1,2^. These profound neurological deficits highlight the essential role of CDKL5 in the normal development and function of the central nervous system. Despite its clinical significance, the pathogenic mechanisms underlying CDD remain largely elusive, and no effective therapeutic interventions are currently available.

Intellectual disability, a core and early-manifesting feature of CDD^2,3^, may arise from altered brain wiring during development or impaired learning and memory processes, potentially due to dysregulation of synaptic plasticity at the cellular level. *Cdkl5* knockout mouse models recapitulate key aspects of CDD, including learning and memory deficits^4,5^, aberrant synaptic activity^6–10^, and structural abnormalities in dendritic spines of cortical neurons, characterized by reduced spine density, increased spine length, and diminished spine head size^7^. While CDKL5 interacts with synaptic proteins and modulates glutamate receptor expression^10,11^, its precise role in synaptic plasticity and the molecular mechanisms involved remain unclear.

CDKL5 functions as a phosphokinase in multiple subcellular compartments through its N-terminal kinase domain, targeting diverse substrates with distinct roles. However, direct evidence linking its phosphorylation activity to synaptic plasticity is limited. The C-terminal domain of CDKL5, less-characterized than its kinase domain, facilitates its localization to excitatory postsynaptic sites via interactions with postsynaptic density protein 95 (PSD95)^12^. Recent studies have revealed that liquid-liquid phase separation (LLPS) drives the assembly and activity-dependent reorganization of the postsynaptic density (PSD)^13,14^, a protein-dense layer beneath the postsynaptic membrane that orchestrates signal transduction and actin cytoskeletal dynamics. PSD95 and other postsynaptic scaffolding proteins undergo LLPS within the PSD through multivalent interactions^13,14^. The size and composition of the PSD are closely correlated with dendritic spine volume and synaptic strength^15,16^. During long-term potentiation (LTP) and long-term depression (LTD), the PSD undergoes extensive structural and compositional reorganization, which is critical for these forms of plasticity^17–19^. As a PSD-enriched component, CDKL5 is likely to play an important role in LLPS-mediated PSD reorganization and synaptic dynamics.

In this study, we demonstrate that CDKL5 undergoes LLPS autonomously in vitro and in cultured neurons. Through its LLPS capability, CDKL5 translocates to excitatory synapses in response to neuronal activity and engages in multivalent interactions with PSD95, promoting the assembly of PSD95-mediated condensates. We further identify Kalirin7, a Rho family guanine nucleotide exchange factor (GEF), as a downstream effector of CDKL5, linking activity-dependent dendritic spine remodeling to actin dynamics. Missense mutations identified in CDD patients disrupt the LLPS capacity of CDKL5 and impair the structural plasticity of synapses, suggesting that dysregulation of LLPS contributes to the pathogenic mechanisms underlying CDD.

## Results

### Activity-dependent translocation of CDKL5 into the PSD promotes spine enlargement

To investigate the dynamic response of CDKL5 to synaptic activity, we chemically induced long-term potentiation (c-LTP) and long-term depression (c-LTD) in cultured hippocampal neurons using glycine and NMDA, respectively. Live-cell imaging and immunostaining consistently demonstrated an activity-dependent redistribution of CDKL5, showing significant enrichment in PSDs following c-LTP induction (Fig. 1a, b, Extended Data Fig. 1a, c) and dispersion from PSDs upon c-LTD stimulation (Fig. 1c, d, Extended Data Fig. 1b, d). These findings indicated that the synaptic localization of CDKL5 is bidirectionally regulated by synaptic activity, being potentiated by c-LTP and attenuated by c-LTD.

**Fig. 1.**
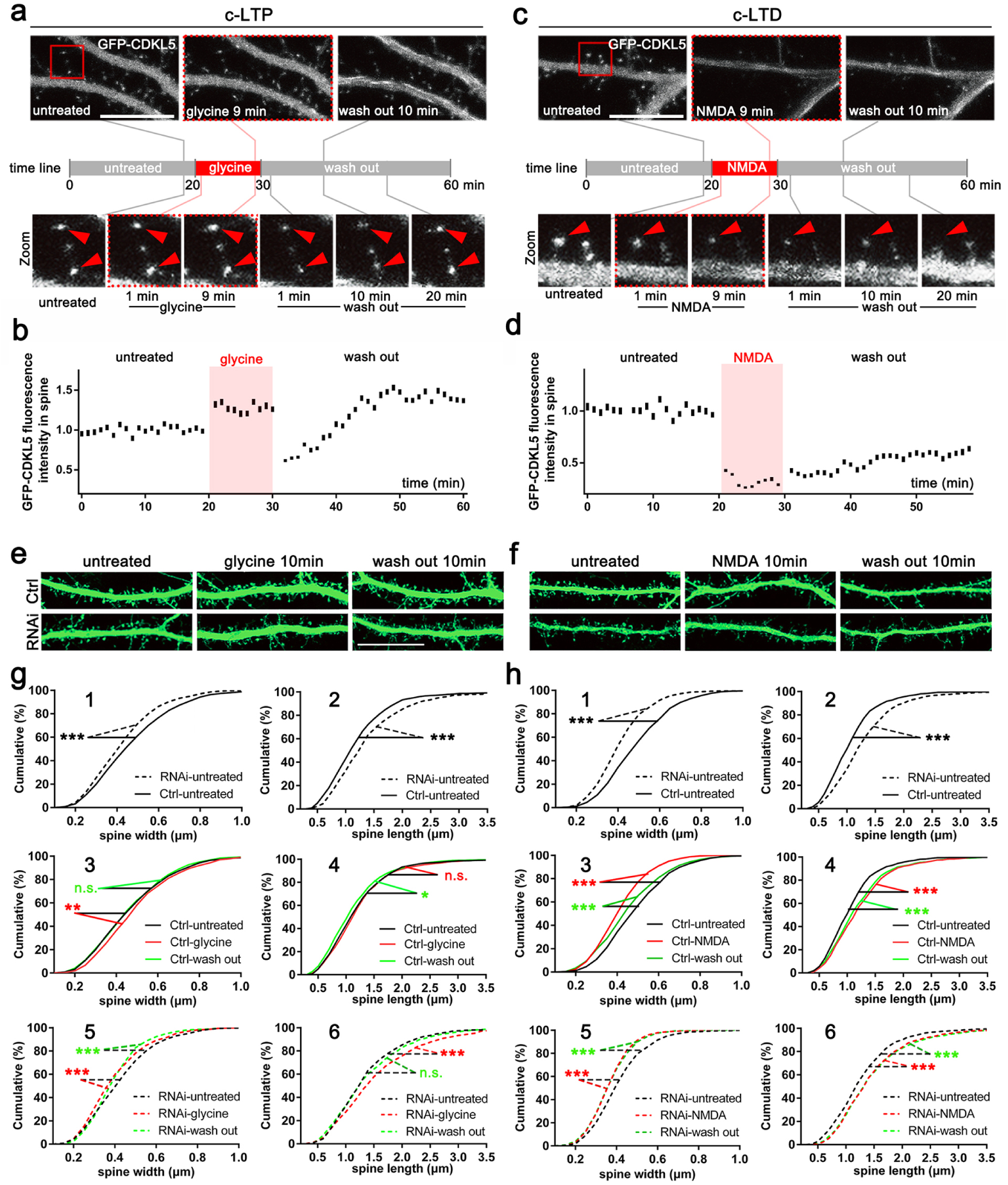
CDKL5 translocation induced by c-LTP and c-LTD is indispensable for structure remodeling of dendritic spines. **a**, Time-lapse images showing glycine-induced increase in GFP-CDKL5 fluorescence intensity in spines. The timeline in the center indicates the sequence of treatments and image capture time points. The red square in the untreated image outlines the region shown in the zoomed-in panel. Red arrowheads in the zoomed-in images highlight spine enlargement following glycine treatment. Scale bar: 20 μm. **b**, Quantification of synaptic GFP-CDKL5 fluorescence intensity during c-LTP in (**a**). A total of 80-120 spines from each time-lapse image were quantified. **c**, Time-lapse images showing NMDA-induced reduction of synaptic GFP-CDKL5 fluorescence intensity. The red square in the untreated image indicates the zoomed-in areas in the lower panel. Red arrowheads in the zoomed-in images indicate spines that underwent shrinkage after NMDA treatment. Scale bar: 20 μm. **d**, Quantification of synaptic GFP-CDKL5 fluorescence intensity during c-LTD in (**c**). A total of 80-120 spines from each time-lapse image were quantified. **e**, **f**, Representative immunocytochemistry (ICC) images showing spine plasticity of hippocampal neurons transfected with GFP and scrambled or CDKL5 shRNA under glycine (**e**) or NMDA treatment (**f**). After the treatment, cells were incubated in ACSF for 10 min (wash out) before being fixed and stained for GFP. Scale bar: 20 μm. **g**, **h**, Cumulative distribution of spine width and length plotted for each condition in (**e**) and (**f**), respectively. A total of 700-1300 spines on 12-15 dendritic branches from 12-15 neurons were analyzed for each condition. The curves were analyzed using asymptotic two-sample Kolmogorov-Smirnov test. *P < 0.05, **P < 0.01, ***P < 0.001.

To examine the role of CDKL5 in activity-dependent dendritic spine remodeling, we conducted shRNA-mediated knockdown experiments followed by c-LTP or c-LTD. The knockdown efficiency of CDKL5-specific shRNA was confirmed through immunostaining of endogenous CDKL5 in cultured neurons transfected with either a scrambled control (Scr) shRNA or the CDKL5-targeting shRNA (Extended Data Fig. 1e). Downregulation of CDKL5 led to a significant reduction in spine size under basal conditions (Fig. 1e, f, untreated group, quantified in Fig. 1g-1, 2, h-1, 2). As anticipated, c-LTP induction in control neurons resulted in spine enlargement (Fig. 1e, glycine-treated ctrl, quantified in Fig. 1g-3, 4), whereas c-LTD induction caused a marked reduction in spine size (Fig. 1f, NMDA-treated ctrl, quantified in Fig. 1h-3, 4). In contrast, in CDKL5-deficient neurons, c-LTP induction failed to increase spine size (Fig. 1e, glycine-treated RNAi, quantified in Fig. 1g-5, 6), and c-LTD induction significantly increased the number of elongated, slender protrusions (Fig. 1f, NMDA-treated RNAi, quantified in Fig. 1h-5, 6).

Collectively, these results suggest that CDKL5 exhibits dynamic translocation into and out of the PSD in response to synaptic activity, and that CDKL5 deficiency disrupts the c-LTP-induced enlargement of dendritic spines, highlighting its critical role in activity-dependent remodeling of synaptic structures.

### CDKL5 undergoes LLPS

To improve the visualization and precise analysis of CDKL5 within cellular compartments, we generated a knock-in mouse line in which a 3x Hemagglutinin (HA) tag was fused to the N-terminus of CDKL5 (Extended Data Fig. 2a). Neurons derived from these mice were subjected to HA immunostaining, allowing for the detection of endogenous CDKL5 with enhanced specificity compared to the CDKL5 antibody. Consistent with prior findings, the majority of CDKL5 was observed to co-localize with PSD95 in dendritic puncta (Fig. 2a). However, CDKL5 also formed distinct, condensed puncta in the soma and nucleus, regions where PSD95 is absent (Fig. 2a-1#; the nucleus region is marked with dotted outline). Notably, within dendrites, certain CDKL5 puncta exhibited no co-localization with PSD95 (Fig. 2a-2#, indicated by white triangles). The droplet-like morphology of CDKL5 resembles the phase-separated PSD condensates formed by PSD95-mediated protein complexes, raising the intriguing possibility that CDKL5 may also possess the capacity for LLPS.

**Fig. 2.**
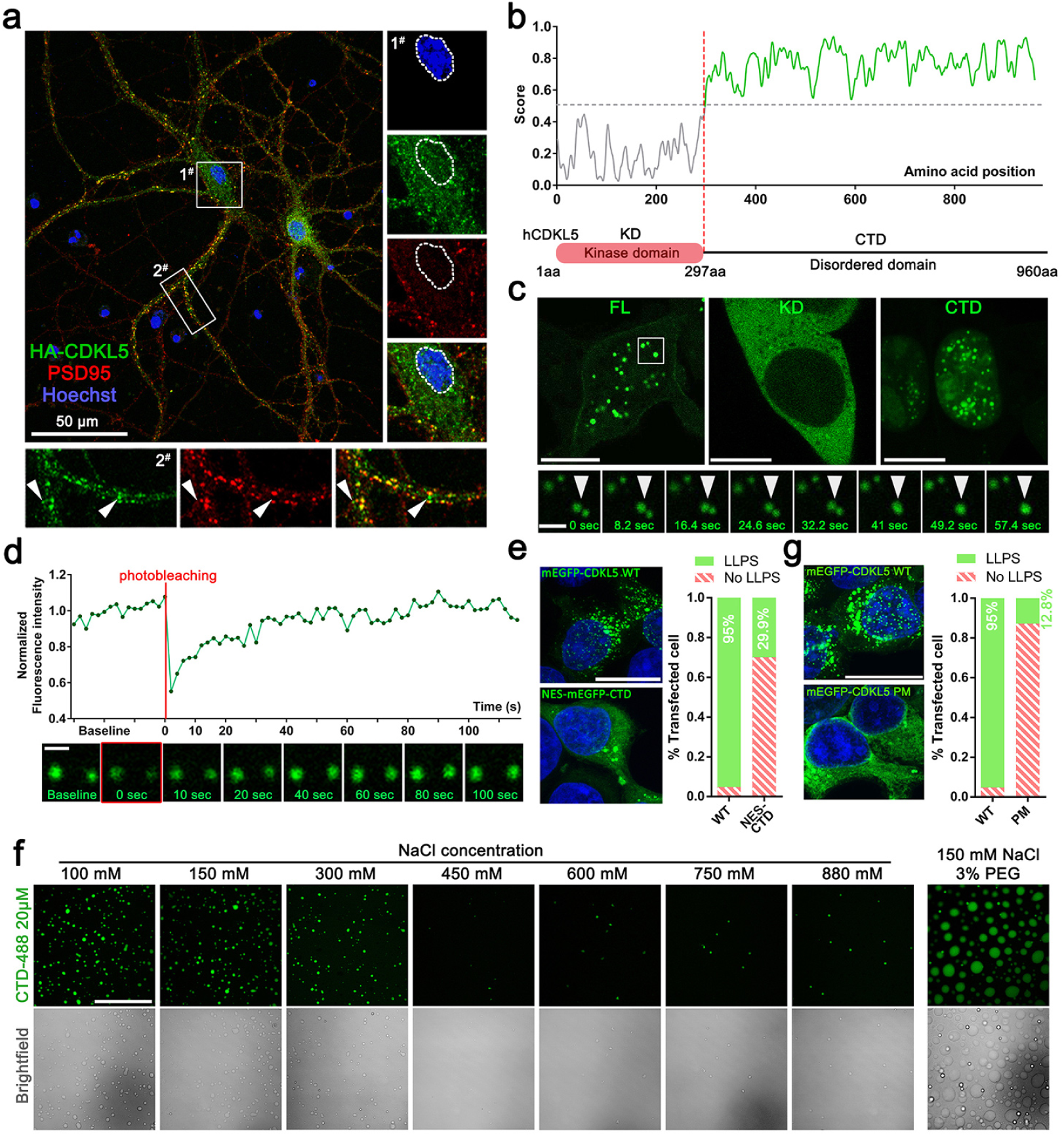
CDKL5 undergoes LLPS through its intrinsically disordered domain, driven by multiple forces. **a**, Representative ICC image of HA-CDKL5 neurons showing the endogenous CDKL5 expression pattern. The white boxes indicate the soma (1^#^) and dendritic regions (2^#^) shown in the zoomed-in images on the right and lower panels, respectively. Arrowheads indicate CDKL5 puncta that did not co-localize with the PSD95 signals. Scale bar: 50 μm. **b**, Intrinsically disordered domain prediction for the human CDKL5 protein sequence using IUPred3 (https://iupred3.elte.hu/). The score for each residue was represented by a continuous line, with a score greater than 0.5 considered disordered. The kinase domain (1-297 aa) was shown in grey, while the C-terminal domain (297-960 aa) was marked in light green. The lower panel shows a protein structure domain diagram for CDKL5. **c**, Representative images (single slices) showing the puncta-formed mEGFP-CDKL5 WT (FL) and CTD, and the diffused distribution of mEGFP-KD when overexpressed in 293T cells. Scale bar: 10 μm. The white box in the FL image highlights magnified time-lapse images that capture two puncta (indicated by white arrowhead) merging into a larger one within one minute. Scale bar: 2 μm. **d**, Representative time-lapse images (bottom) and quantification (top) of mEGFP-hCDKL5 puncta intensity in 293T cells from the fluorescence recovery after photobleaching (FRAP) experiment at the indicated time points. Scale bar: 2 μm. **e**, Images and quantification of the percentage of LLPS cells transfected with a nuclear export signal (NES) tagged CTD (NES-mEGFP-CTD) or mEGFP-CDKL5-FL. CTD was found to mainly localize in the nucleus; to make a fair comparison with FL, an NES was employed to relocate CTD to the cytoplasm. Scale bar: 20 μm. **f**, Images of protein droplets formed by 20 μM of CDKL5 298-960 aa protein labeled with BDP FL dye (emission collected at 488 nm) under various buffer conditions. Scale bar: 50 μm. **g**, Images and quantification of the percentage of LLPS cells transfected with mEGFP-CDKL5 WT or the PM mutation (using the same WT data as in (**e**) to calculate the percentage of LLPS cells). Scale bar: 20 μm.

The CDKL5 protein comprises an N-terminal phosphokinase domain and an extended C-terminal region. Structural predictions reveal that, aside from the kinase domain (CDKL5^KD^, residues 1-297), the extended C-terminal region (CDKL5^CTD^, residues 297-960) constitutes an intrinsically disordered region (IDR) that is fully disordered^20,21^ (Fig. 2b), a feature frequently associated with the capacity for phase separation^22,23^. To investigate whether CDKL5 can undergo LLPS, we transfected monomeric enhanced green fluorescent protein (mEGFP) tagged full-length CDKL5 (CDKL5^FL^), CDKL5^KD^, and CDKL5^CTD^ into human embryonic kidney (HEK) 293T cells. We observed that both CDKL5^FL^ and CDKL5^CTD^ formed bright and spherical puncta (Fig. 2c), whereas CDKL5^KD^ displayed a diffuse cytoplasmic distribution (Fig. 2c). The CDKL5^FL^ puncta exhibited fluid-like properties, as evidenced by the fusion of two puncta into a larger one (Fig. 2c, bottom row). Furthermore, mEGFP-CDKL5^FL^ puncta showed fluorescence recovery after photobleaching (Fig. 2d), indicating dynamic exchange of components with the surrounding dilute phase. Although CDKL5^CTD^ retained the ability to undergo LLPS, it displayed a significantly reduced propensity for puncta formation compared to CDKL5^FL^ (Fig. 2e). Given that CDKL5^FL^ can recruit CDKL5^KD^ into its puncta, while CDKL5^CTD^ lacked this capability (Extended Data Fig. 2b), we proposed that the diminished LLPS efficiency of CDKL5^CTD^ may result from the absence of additional binding valency mediated by KD-KD homo-interactions. Collectively, these data suggest that both the KD and CTD contribute to the complete and efficient LLPS of CDKL5.

To rule out the possibility that CDKL5 droplet formation in cells results from coalescence with other protein condensates or depends on interactions with additional proteins, we aimed to purify the CDKL5 protein and examine its LLPS capability in vitro. Unfortunately, due to technical challenges, we were unable to obtain qualified full-length CDKL5 protein, but we successfully purified the CDKL5^CTD^ protein instead. Under conditions of pH 7.4, 150 mM NaCl, and room temperature, fluorescently labeled CDKL5^CTD^ formed spherical droplets at concentrations of 3 µM or higher, demonstrating that CDKL5 possesses an intrinsic capability for LLPS independent of external factors (Extended Data Fig. 2c). Notably, the number and size of CDKL5^CTD^ droplets decreased progressively with increasing NaCl concentrations ranging from 100 mM to 880 mM (Fig. 2f), whereas the addition of the crowding agent PEG8000 enhanced CDKL5^CTD^ LLPS (Fig. 2f). Taken together, these findings suggest that CDKL5 can autonomously undergo LLPS.

LLPS is driven by various types of multivalent intermolecular interactions, including charge-charge, cation-π, π-π, hydrophobic, and hydrogen-bonding interactions^24^. To explore how amino acid composition governs the physical properties of CDKL5^CTD^ LLPS, we analyzed the sequence of its IDR. We observed that CDKL5^CTD^ contains both positively and negatively charged residues (Extended Data Fig. 2d), which could potentially facilitate charge-charge interactions critical for LLPS. However, selective alanine substitution of charged residues at various positions within the CTD did not disrupt phase separation (Extended Data Fig. 2e). Moreover, we noted that CDKL5^CTD^ comprises 32 aromatic amino acid residues evenly distributed across the CTD (Extended Data Fig. 2d), which are likely to mediate cation-π and π-π interactions indispensable for LLPS. Notably, partial substitution of specific aromatic residues significantly impaired CDKL5 LLPS (Extended Data Fig. 2f), while mutation of all 32 aromatic residues (phenylalanine to alanine, tyrosine to serine, and tryptophan to leucine), designated as Phenyl groups Mutated (CDKL5^PM^), nearly abolished LLPS (Fig. 2g). These results suggest that in the CTD, the amino acid residues contributing to CDKL5 LLPS are uniformly distributed, with aromatic residues playing a predominant role in driving this process.

### CDKL5 forms co-condensates with PSD95

Based on the PSD localization and LLPS properties of CDKL5, we hypothesized that CDKL5 participates in the assembly of PSD-associated condensates through co-condensation with PSD95. To test this hypothesis, we conducted *in vitro* reconstitution assays by mixing purified CDKL5^CTD^ with full-length PSD95 protein at varying stoichiometric ratios. Phase-contrast microscopy revealed the formation of heterotypic condensates containing both proteins (Fig. 3a). Importantly, the addition of PSD95 lowered the threshold concentration required for CDKL5^CTD^ to undergo LLPS, and dramatically increased the size of CDKL5^CTD^ droplets (Fig. 3a), indicating that PSD95 promotes the LLPS of CDKL5. Quantitative analysis showed that these co-condensates formed across a wide range of stoichiometric ratios (CDKL5:PSD95 = 1:50 to 4:1) and protein concentrations (1.5-24 µM at a 1:5 molar ratio), indicating robust co-condensation between CDKL5 and PSD95. Notably, the formation of the condensed co-droplets was readily observed at both the physiologically relevant stoichiometric range, as determined based on immunoblotting analysis of tissue lysates from adult mouse brain (PSD95: CDKL5 ≈ 10:1, as determined in Extended Data Fig. 3a), and biologically relevant protein concentration (∼ 1 μM of PSD95 or lower) (Fig. 3a, the physiologically relevant stoichiometric range was indicated with a red dashed box), suggesting that the CDKL5-PSD95 co-condensates can form under biologically meaningful conditions.

**Fig. 3.**
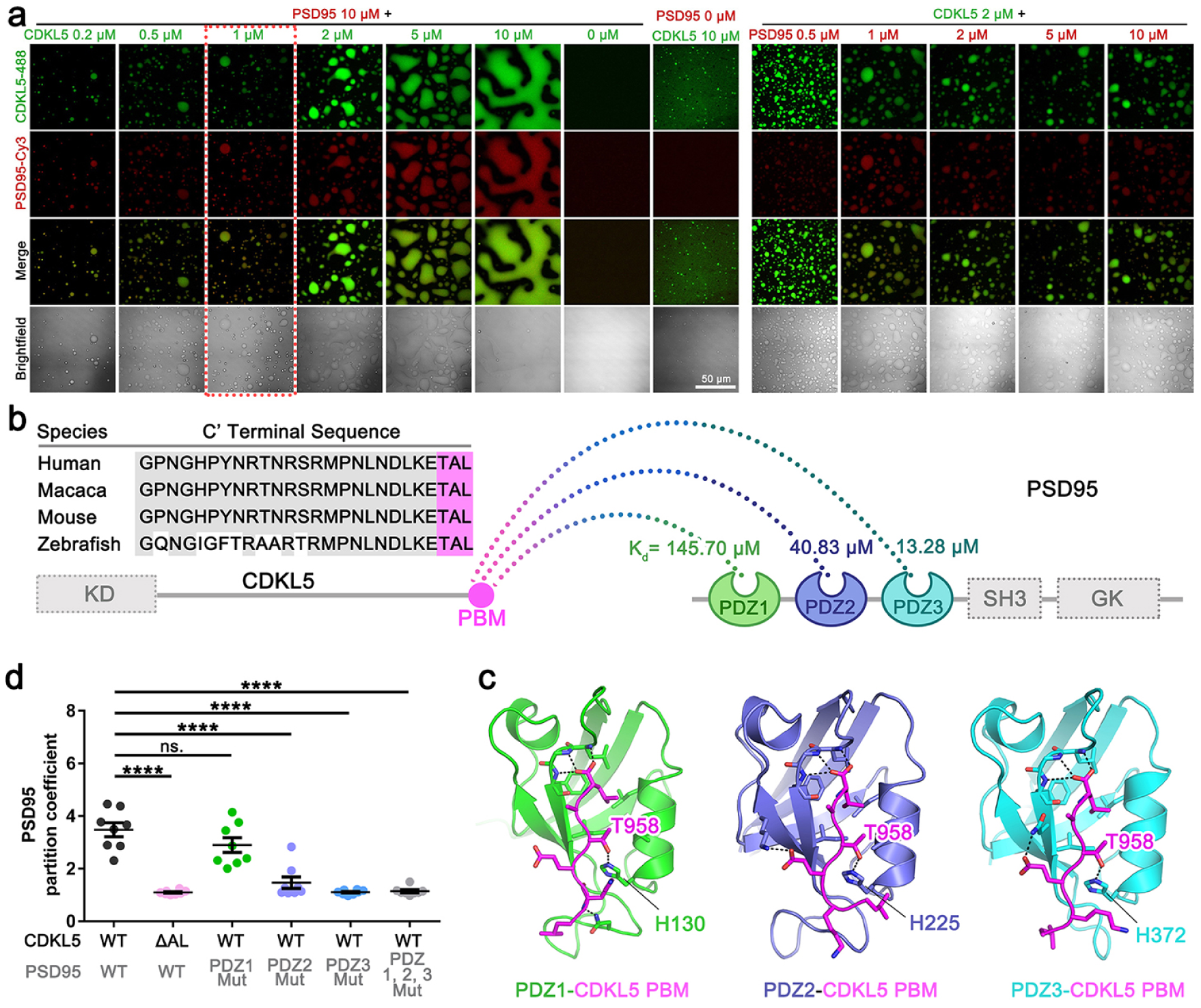
The interaction of CDKL5 with the PDZ domains of PSD95 contributes to the formation of CDKL5-PSD95 co-condensates and its PSD localization. **a**, Images showing droplets formed in vitro by CDKL5-PSD95 co-condensate at different CDKL5:PSD95 molar ratios. The brightfield images are single slices taken at the optimal focal distance and adjusted separately. Scale bar: 50 μm. **b**, Alignment of the C-terminal end sequence of the CDKL5 protein across different species. The conserved core sequences TAL for the PBM are highlighted with a carmine background. The binding targets for the PBM of CDKL5, the three PDZ domains of PSD95, are illustrated in a diagram marked with the *K_d_* calculated from fluorescence polarization-based assays. **c,** Predicted structures of complexes formed by the PBM of CDKL5 and the PDZ domains of PSD95 using the AlphaFold3 server (https://alphafoldserver.com/). **d**, Quantification of partition coefficients of PSD95 in cells co-expressing PSD95 WT with CDKL5 WT or ΔAL, or PSD95 PDZ mutations with CDKL5 WT. For the PSD95 PDZ mutations, one or all PDZ domains were inactivated through single amino acid mutations, as shown in Extended Data Fig. (**4c**). Eight individual cells from each group were analyzed using unpaired two-tailed t-test. ****P < 0.0001. Data are represented as mean ± SEM.

Next, we sought to identify the essential factor governing CDKL5-PSD95 co-condensates formation. The PDZ domains (named after the three proteins: postsynaptic density 95, **PSD95**; discs large, **D**lg; zonula occludens-1, **Z**O-1)^25^ of PSD95 are known to facilitate multivalent interactions with other scaffolding proteins, thereby driving the formation of PSD condensates. Meanwhile, we noticed that the last three amino acid residues of the human CDKL5 (-TAL) potentially constitute an evolutionarily conserved PDZ-binding motif (PBM) that is capable of interacting with class I PDZ domains (Fig. 3b). Indeed, fluorescence polarization-based assays demonstrated that CDKL5^PBM^ binds to the PDZ1, PDZ2, and PDZ3 domains of PSD95 with dissociation constants (*K*_d_) of ∼ 145.7 μM, 40.83 μM, and 13.28 μM, respectively (Fig. 3b and Extended Data Fig. 3b). Moreover, we employed AlphaFold3 to generate structural models for the PDZ-PBM complexes, revealing that CDKL5^PBM^ interacts with PSD95 PDZ domains in a typical PDZ-PBM binding mode. For example, the side chain of CDKL5 T958 interacts with the imidazole rings of H130^PDZ1^, H225^PDZ2^, and H372^PDZ3^ of PSD95, respectively (Fig. 3c). These results provided compelling evidence that CDKL5 engages with PSD95 through multivalent interactions through its PDZ domains.

To investigate whether PBM-PDZ interactions between CDKL5 and PSD95 contribute to co-condensation formation, we introduced point mutations into each PDZ domain of PSD95 (PSD95^PDZ1mut^: PSD95^H130A^, PSD95^PDZ2mut^: PSD95^H225A^, PSD95^PDZ3mut^: PSD95^H372A^ and triple mutant PSD95^PDZ123mut^: H130A/H225A/H372A) to selectively disrupt their PBM binding capacity. Co-expression analysis revealed that PSD95^PDZ2mut^, PSD95^PDZ3mut^, and PSD95^PDZ123mut^ lost their ability to form co-condensates with CDKL5 compared to wild-type PSD95 (Fig. 3d and Extended Data Fig. 3c), whereas PSD95^PDZ1mut^ exhibited a relatively minor effect on co-condensate formation. These results established that the CDKL5 PBM preferentially interacts with the PDZ2 and PDZ3 domains of PSD95. Notably, deletion of the last two amino acid residues of CDKL5 (referred to as CDKL5^ΔAL^), resulted in a reduction in co-condensation comparable to that observed with the PSD95^PDZ123mut^ triple mutant, as quantified by partition coefficient analysis (Fig. 3d, Extended Data Fig. 3c). Furthermore, structural analysis demonstrated that T958 of CDKL5 is crucial for its binding to the PDZ domains of PSD95 (Fig. 3c). Consistent with this, the CDKL5^T958R^ mutation, identified in a CDD patient^26^ showed a significantly reduced binding affinity for the PDZ2 and PDZ3 domains of PSD95 (Extended Data Fig. 3d).

Together, these findings demonstrate that the multivalent PBM-PDZ interactions between CDKL5 and PSD95 drive co-condensates formation, and impaired PBM-PDZ interactions result in reduced PSD95 enrichment within CDKL5 puncta, reflecting defects in CDKL5-PSD95 co-condensate formation.

### Formation of CDKL5-PSD95 co-condensates is essential for synaptic targeting of CDKL5 and the maintenance of PSD architecture

We next set out to explore the impact of CDKL5 LLPS and its co-condensation with PSD95 on the synaptic targeting of CDKL5. To this end, we took advantage of a mouse model carrying a truncation mutation of CDKL5 that results in a completely absence of the CDKL5 protein^27^ (referred to as 493-stop mice, Extended Data Fig. 4a). We then analyzed the spine-to-shaft fluorescence intensity ratio of various CDKL5 mutants expressed in neurons derived from 493-stop mice, which lack endogenous CDKL5. Our analysis revealed that CDKL5^CTD^ maintained substantial colocalization with PSD95 within the PSD (Fig. 4a, b), albeit with a significantly reduced efficiency compared to wild-type CDKL5 (Fig. 4a, b), probably due to the incomplete LLPS resulting from the deletion of the KD (Fig. 2). In contrast to CDKL5^CTD^, CDKL5^KD^ exhibited predominant dendritic shaft localization (Fig. 4a, b). To figure out whether the LLPS capacity and the PBM-mediated CDKL5-PSD95 interaction are both required for the synaptic localization of CDKL5, we analyzed PSD enrichment of two distinct mutants: the phase separation-deficient mutant CDKL5^PM^, which retains an intact PBM but is incapable of LLPS (Extended Data Fig. 4b), and the PBM deletion mutant CDKL5^ΔAL^, which preserves LLPS capacity. Quantitative analysis demonstrated a complete loss of synaptic localization for the CDKL5^PM^ mutant, with a predominant distribution in dendritic shafts (Fig. 4a, b), whereas the PBM-deleting mutant (CDKL5^ΔAL^) showed significant retention in PSDs, although it was reduced compared to WT (Fig. 4c). Consistent with the CDKL5^ΔAL^ variant, the CDD-associated CDKL5^T958R^ mutant, which disrupts PBM, exhibited impaired synaptic targeting (Fig. 4d). Moreover, we found that the CDKL5 kinase-dead mutation (K42R)^28^ did not affect the LLPS propensity of CDKL5 (Extended Data Fig. 4c), the formation of CDKL5-PSD95 co-condensates (Extended Data Fig. 4d), or synaptic distribution pattern (Extended Data Fig. 4e).

**Fig. 4.**
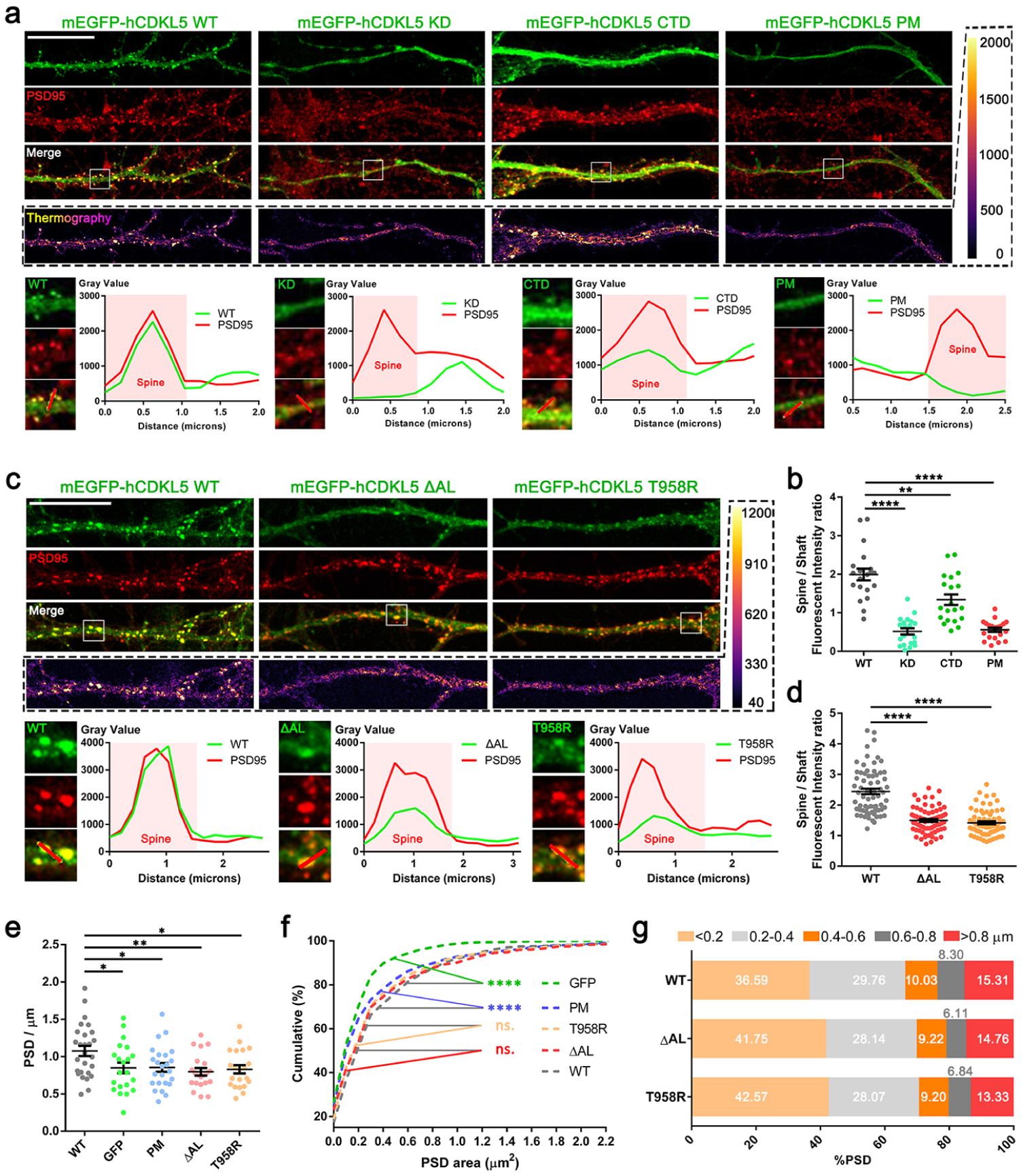
The PBM-PDZ interaction between CDKL5 and PSD95, along with the LLPS of CDKL5, is essential for CDKL5 localization at the PSD. **a**, Images of CDKL5-deficient neurons (493-stop neurons) expressing mEGFP-CDKL5 WT, KD, CTD, and PM. CDKL5-PSD95 colocalization is presented as thermographic images in the bottom row. The white boxes in the merged images indicate the ROIs (regions of interest) where spines were selected for line plotting performed in the lower panel. PSD95 was used as a marker to define the spine region, and the average intensity in the spine area and shaft area was utilized to calculate the spine-shaft ratio. Scale bar: 20 μm. **b**, The spine-shaft intensity ratio from (**a**) is calculated and presented as mean ± SEM. Each group includes data from 10 spines from 2 neurons. Unpaired t-test with two tails was used, **P < 0.01, ****P < 0.0001. **c**, Fluorescence images of CDKL5 and PSD95 from CDKL5-deficient (493-stop) neurons expressing mEGFP-CDKL5 WT, PM, or T958R. CDKL5-PSD95 colocalization is presented as thermographic images in the bottom row. The white boxes in the merged images indicate the spines selected for line plotting performed in the lower panel. Scale bar: 10 μm. **d**, The Spine-shaft intensity ratio for each group shown in (**c**), each group includes data of 70 spines from 7 neurons. Mann-Whitney U test with two tails was used, ****P < 0.0001. Data are represented as mean ± SEM. **e**, Quantifications of PSD density for CDKL5-deficient neurons expressing mEGFP-CDKL5 WT, PM, ΔAL, or T958R, GFP expressed by lentivirus served as negative control. Each dot represents an individual dendrite, with sample sizes of n=27 (WT), 21 (GFP), 24 (PM), 21 (ΔAL), and 21 (T958R). Unpaired two-tailed t-test was used for the analysis. *P < 0.05, **P < 0.01. Data are represented as mean ± SEM. **f**, The cumulative distribution of PSD area for each condition in (**e**). The sample sizes were n=657 for GFP, 1161 for WT, 895 for PM, 869 for ΔAL, and 848 for T958R. The curves were analyzed using asymptotic two-sample Kolmogorov-Smirnov test, with PSD puncta larger than 5 μm² excluded as multiple overlay spines. ****P < 0.0001. **g**, PSD area size frequency distribution of WT, ΔAL and T958R, plotted with data from (**f**).

Next, we sought to explore the impact of these mutations on PSD morphology by quantifying the area of PSD95-positive puncta, a well-established morphological parameter that reflects PSD dimensions and strongly correlates with spine size. We performed this experiment in CDKL5-depleted neurons, in which CDKL5 WT, its mutants, or GFP (the negative control) were exogenously expressed so that their functional effects could be analyzed and compared. We found that the expression of CDKL5 WT significantly increased PSD density and size compared to that of GFP (Fig. 4e, f). However, CDKL5^PM^, CDKL5^ΔAL^, and CDKL5^T958R^ all exhibited compromised effects on scaling up the PSD puncta area compared to WT (Fig. 4e) (Fig. 4f and 4g). Furthermore, quantitative analyses revealed that CDKL5 fluorescence intensity was strongly correlated with both PSD95 fluorescence intensity and PSD puncta area (Extended Data Fig. 4f, g), suggesting that CDKL5 promotes PSD assembly in a dose-dependent manner.

These data collectively indicate that while kinase activity is dispensable, both the direct interaction and co-condensation between CDKL5 and PSD95 are essential for the synaptic recruitment of CDKL5 and the maintenance of normal PSD size.

### Activity-dependent synaptic translocation of CDKL5 requires co-condensation of CDKL5 and PSD95

Next, we explored whether the formation of the CDKL5-PSD95 co-condensates is required for the activity-dependent translocation of CDKL5 into the PSD. CDKL5-depleted neurons expressing mEGFP-CDKL5 WT, PM, ΔAL, and T958R via lentiviral infection were subjected to c-LTP or c-LTD stimulation, followed by immunostaining to analyze the co-localization of CDKL5 and PSD95. Consistent with the patterns of endogenous CDKL5 (Extended Fig. 1a, b), mEGFP-CDKL5 WT displayed an increased colocalization with PSD95 in synapses following c-LTP and a decreased colocalization after c-LTD (Fig. 5a). CDKL5^PM^ exhibited diminished colocalization with PSD95 under baseline conditions (Fig. 5b), and failed to change its colocalization pattern with PSD95 in response to c-LTP or c-LTD stimulation (Fig. 5b). In contrast, both the CDKL5^ΔAL^ and CDKL5^T958R^ mutants showed a notable decrease in synaptic localization during c-LTP; with the reduction being even more pronounced during c-LTD (Fig. 5c, d). These data demonstrate that both LLPS of CDKL5 and its co-condensation with PSD95 are crucial for the activity-dependent translocation of CDKL5 into and out of the PSD.

**Fig. 5.**
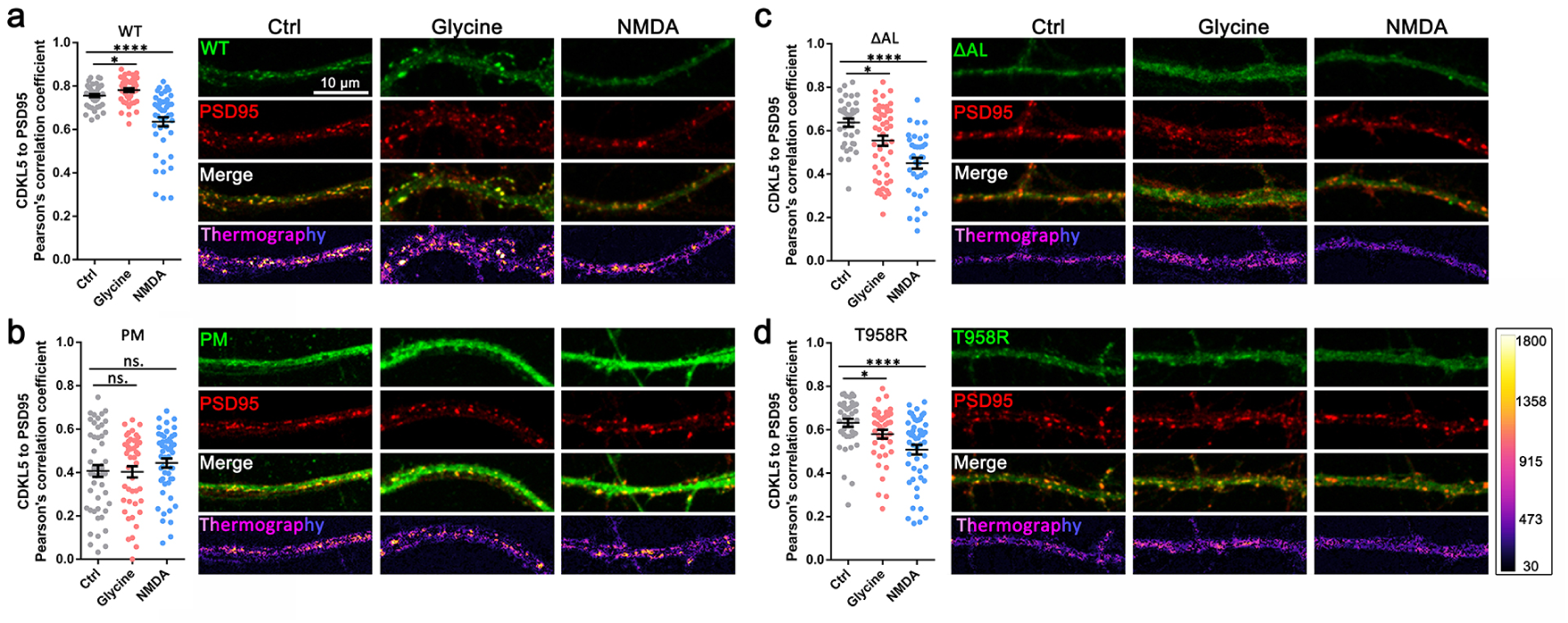
The formation of CDKL5-PSD95 co-condensates is required for activity dependent trafficking of CDKL5 into the PSD. **a, b, c, d,** Pearson’s correlation coefficient plots of CDKL5 and PSD95, along with fluorescence images from CDKL5-deficient (493-stop) neurons expressing mEGFP-CDKL5 WT (**a**), PM (**b**), ΔAL (**c**), or T958R (**d**) under untreated, c-LTP or c-LTD stimulation. CDKL5-PSD95 colocalization is presented as thermographic images in the bottom row. For the control, c-LTP (Glycine), and c-LTD (NMDA) conditions in each group, the sample sizes are as follows: 45, 45, 45 for WT; 35, 50, 35 for ΔAL; 40, 40, 50 for T958R; and 50, 45, 50 for PM. Two-tailed Mann-Whitney U test was used for statistical analysis. *P < 0.05, ****P < 0.0001. Data are represented as mean ± SEM. Scale bar: 10 μm.

### The CDKL5-PSD95 co-condensates recruit Kalirin7 RhoGEF into the PSD

Although the increasing enrichment of CDKL5 within the PSD coincides with spine growth during LTP, such PSD rearrangement alone is not sufficient to elicit comprehensive alterations in spine morphology. The enlargement of spines induced by LTP also requires the assembly of the actin cytoskeleton^29,30^. Given that CDKL5^PM^ failed to induce activity-dependent synaptic enrichment and spine enlargement, we hypothesized that LLPS of CDKL5 is critical for the recruitment of the downstream signaling modulator(s) involved in the activity-dependent remodeling of the cytoskeleton. To identify potential targets, we conducted an assay using the phase separated CDKL5^CTD^ as a bait to isolate candidates from mouse brain lysates (Fig. 6a). After electrophoresis, the gel segment containing the CTD condensate-enriched proteins (Fig. 6b, indicated by the red pyramid) was subjected to mass spectrometric analysis. Among the approximately 50 candidates enriched in the CTD protein condensate, we selected Kalirin7 for further characterization because it ranks in the top 3 on the list and is abundantly present in synapses. Its presence in the CTD protein condensate was validated by immunoblotting analysis (Fig. 6b, right panel).

**Fig. 6.**
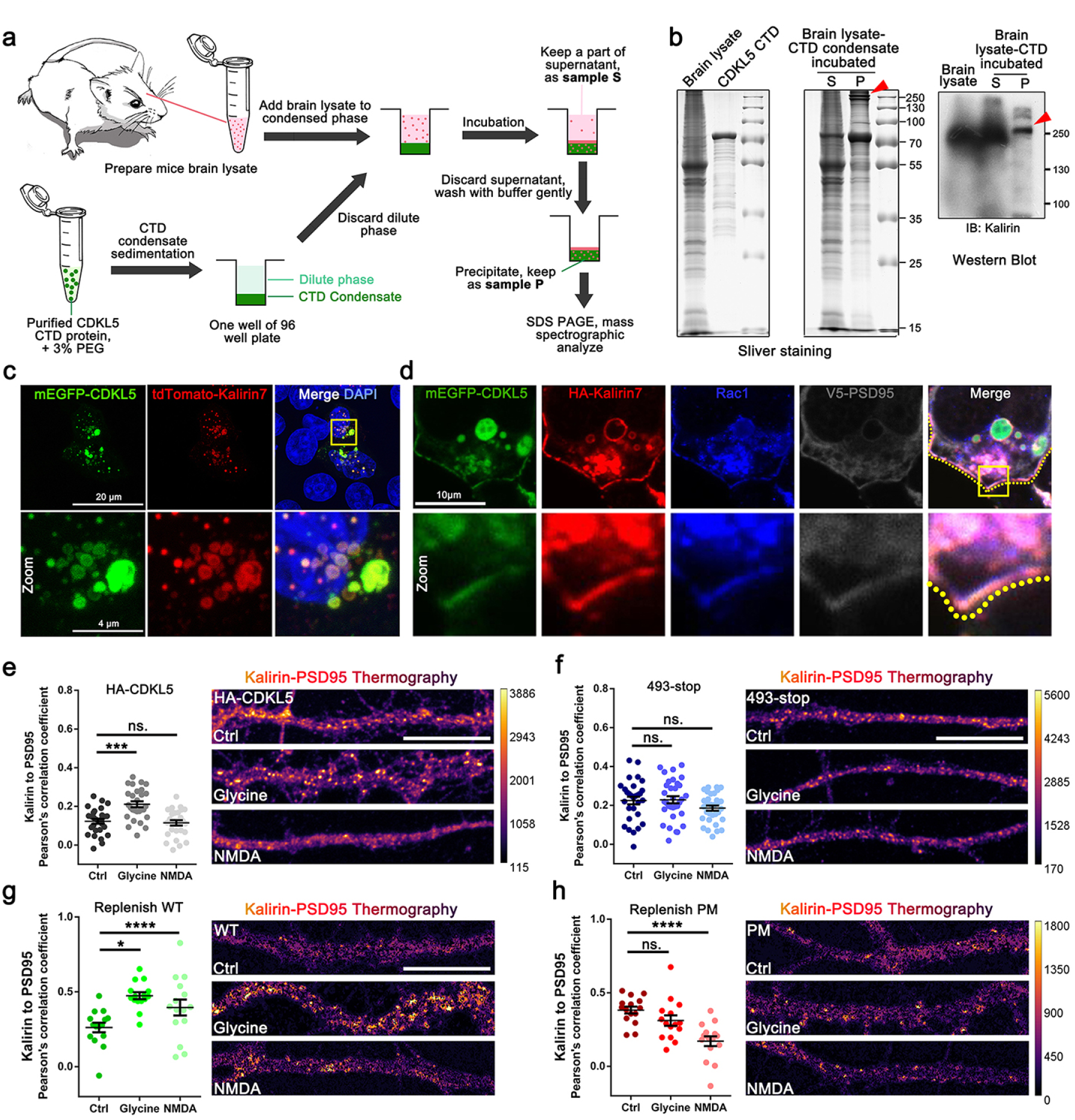
CDKL5 recruits Kalirin7 into its condensates and is responsible for the activity-dependent recruitment of Kalirin7 into the PSD. **a**, Experimental scheme for CDKL5^CTD^ condensate pull-down assay. **b**, Silver staining results (left panel) of the supernatant (S) and pellet (P) fractions from the CDKL5^CTD^ pull-down assay, and western blot validation of Kalirin (right panel). The red arrowhead in the left panel indicates potential interaction proteins of the CDKL5 condensate. **c**, Representative images of 293T cells co-expressing mEGFP-CDKL5 and tdTomato-Kalirin7. Scale bar: 20 μm. The yellow box indicates zoomed-in regions shown in the lower panel. Scale bar: 4 μm. **d**, Representative images of 293T cells co-expressing mEGFP-CDKL5, HA-Kalirin7, and V5-PSD95. Endogenous RAC1 was labeled in blue. Scale bar: 10 μm. The dotted yellow line denotes the outline of the cell, and the yellow box indicates zoomed-in regions shown in the lower panel. **e**, **f**, Pearson’s correlation coefficient and thermography for Kalirin-PSD95 colocalization in c-LTP and c-LTD-stimulated HA-CDKL5 neurons (e) and 493-stop neurons (f). Scale bar: 20 μm. In the HA-CDKL5 group, a total of 28 dendrite segments from 7 neurons were analyzed. For the 493-stop neurons, 29, 33, and 32 dendrite segments from 7, 8, and 8 neurons in the Ctrl, Glycine, and NMDA groups were analyzed, respectively. Unpaired t-test with two tails was used. ***P < 0.001. Data are represented as mean ± SEM. **g, h**, Pearson’s correlation coefficient and thermography for Kalirin-PSD95 colocalization in c-LTP and c-LTD-stimulated 493-stop neurons expressing CDKL5 WT (g) or PM (h). Both panels share the same calibration bar presented in (**h**). Scale bar: 20 μm. A total of 15 dendrite segments from 3 neurons for each group were analyzed. Unpaired t-test with two tails was used. *P < 0.05, ****P < 0.0001. Data are represented as mean ± SEM.

Kalirin7 is a RhoGEF for Rac1 (Ras-related C3 botulinum toxin substrate 1), which binds to PSD95 to regulate synapse formation and synaptic plasticity^31–33^. During LTP, activation of the NMDA receptor triggers the phosphorylation of Kalirin7 by calcium/calmodulin-dependent protein kinase II (CaMKII), thereby enhancing its GEF activity towards Rac1. The activated Rac1 subsequently drives synaptic actin cytoskeletal dynamics through enhanced polymerization, leading to the rapid enlargement of dendritic spines^34,35^ . Notably, CDKL5 is implicated in orchestrating neuronal morphogenesis through its involvement in the Rac1 signaling pathway^36^. Thus, we thought that Kalirin7 might be a downstream target of the CDKL5-PSD95 co-condensates that can regulate cytoskeletal dynamics during LTP-induced spine enlargement. To test this, we first confirmed the enrichment of Kalirin7 in the CDKL5 condensate. Co-expression of tdTomato-Kalirin7 and mEGFP-CDKL5 showed overlapping signals in HEK 293T cells (Fig. 6c). When co-expressed with CDKL5 and PSD95, Kalirin7 was found to be enriched in the CDKL5-PSD95 co-condensate along with the endogenous Rac1 at the cell periphery, suggesting the formation of CDKL5-PSD95-Kalirin7 triple-component co-condensates (Fig. 6d).

It has been reported that the localization of Kalirin7 in the PSD increased in an activity-dependent manner^34^. We next investigated whether CDKL5 regulates the synaptic localization of Kalirin7 in response to synaptic activity. In WT neurons, c-LTP stimulation elicited a significant augmentation of the colocalization of Kalirin7 and PSD95 within the PSD, while c-LTD stimulation didn’t significantly alter Kalirin-PSD95 co-localization (Fig. 6e). Furthermore, the synaptic translocation of Kalirin7 was strongly correlated with CDKL5, as evidenced by a concurrent increase in the colocalization of Kalirin7 and CDKL5 during c-LTP (Extended Data Fig. 5a, b), suggesting a functional interplay between CDKL5 and Kalirin7 during synaptic plasticity. Importantly, the postsynaptic targeting of Kalirin7 remained unchanged following c-LTP stimulation in the absence of CDKL5 (Fig. 6f and Extended Data Fig. 5c), and was successfully rescued by re-expressing CDKL5 WT but not the LLPS-deficient CDKL5^PM^ mutant (Fig. 6g, h and Extended Data Fig. 5d-f). These data indicate that LLPS of CDKL5 is crucial for the activity-dependent recruitment of Kalirin7 to regulate postsynaptic structural plasticity.

### Disease-associated mutations in CDKL5 impairs its LLPS capacity and synaptic function

Given that CDKL5^PM^, CDKL5^ΔAL^, and CDKL5^T958R^ disrupt CDKL5-PSD95 co-condensates formation and activity-dependent translocation of CDKL5, other CDD-associated mutations might similarly dysregulate synaptic plasticity through this mechanism. To investigate this, we screened CDD-associated single-residue missense mutations instead of truncation mutations, as these are less likely to result in a complete loss of the CDKL5 protein^27^. We selected the previously unreported p.C30W mutation and the p.C291Y mutation^37^ for further analysis. Compared to mEGFP-CDKL5^WT^, mEGFP-CDKL5^C291Y^ exhibited a significant reduction in LLPS in HEK 293T cells (Fig. 7a). In contrast, mEGFP-CDKL5^C30W^ retained LLPS capability but formed significantly smaller puncta with increased density (Fig. 7b-d). Furthermore, the fluorescence recovery rate after photobleaching of mEGFP-CDKL5^C30W^ was markedly slower than that of mEGFP-CDKL5^WT^ (Fig. 7e), suggesting that the CDKL5^C30W^ condensate exhibits higher density and reduced dynamics.

**Fig. 7.**
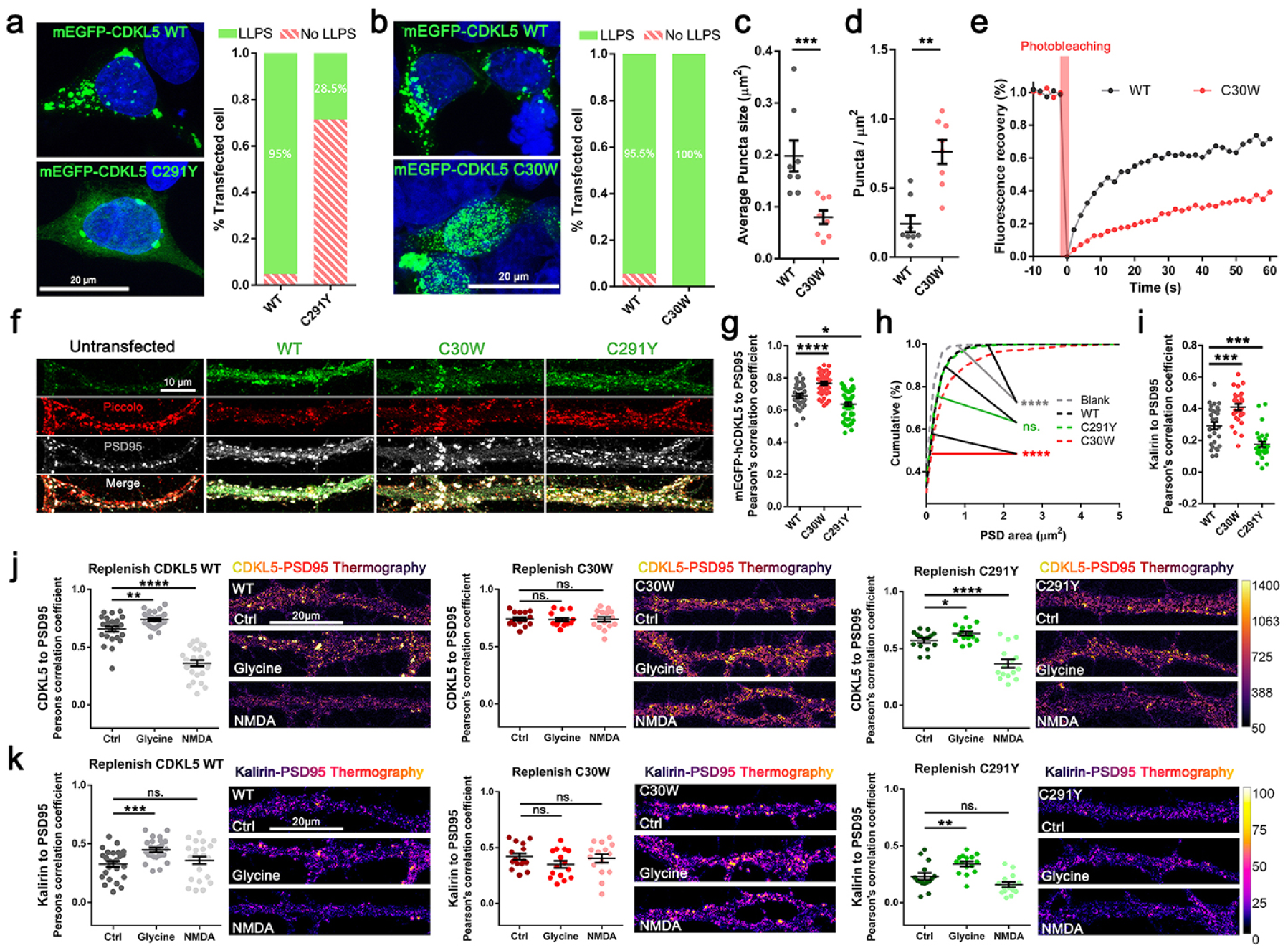
CDD-associated mutations impair LLPS and activity-dependent spine plasticity. **a**, Images and quantification of the percentage of LLPS cells expressing mEGFP-CDKL5 WT or C291Y. Data for C291Y were collected alongside Fig. (**2e**), Fig. (**2g**), and Extended Data Fig.(**4c**), all sharing the same WT data. **b**, Images and quantification of percentage of LLPS cells expressing mEGFP-CDKL5 WT or C30W mutation. **c**,**d**, Quantifications of average puncta size (**c**) and density (**d**) from (**b**) showing that C30W puncta were significantly smaller and denser than those of WT. Each dot represents data from one cell, sample sizes were 8 both for C30W and WT, Mann-Whitney U test with two tails was used. **P<0.01, ***P < 0.001. Data are represented as mean ± SEM. **e**, Quantitative results for FRAP analysis of WT and C30W. A total of 16 and 11 cells were analyzed for WT and C30W, respectively. **f**, Representative images of CDKL5-deficient (493-stop) neurons transfected with lentivirus expressing CDKL5-WT, C30W, or C291Y. Untransfected CDKL5-deficient (493-stop) neurons were used as a negative control. Piccolo and PSD95 were used to define the spine region. **g**, Pearson’s correlation coefficient of CDKL5 to PSD95 in (**f**). A total of 30 branches from 6 neurons, 45 branches from 9 neurons, and 50 branches from 10 neurons were analyzed for WT, C30W, and C291Y, respectively. Mann-Whitney U test with two tails was used. *P < 0.05, ****P < 0.0001. Data are represented as mean ± SEM. **h**, Plots of the cumulative distribution for the PSD area of CDKL5-deficient (493-stop) neurons expressing CDKL5 WT, C30W, or C291Y. Sample sizes were 543 for blank, 731 for WT, 913 for C30W, and 724 for C291Y. PSD puncta larger than 5 μm² were excluded from this test as multiple overlaid spines. The curves were analyzed using asymptotic two-sample Kolmogorov-Smirnov test. ****P < 0.0001. **i,** Comparison of Kalirin to PSD95 Pearson’s correlation coefficient among WT, C30W, and C291Y at basal conditions. A total of 28 dendritic branches from 4 neurons were analyzed for each group. The Mann-Whitney U test with two tails was used. ***P < 0.001. Data are represented as mean ± SEM. **j**, Thermographs and Pearson’s correlation coefficient of CDKL5 to PSD95 in CDKL5-deficient (493-stop) neurons replenished with CDKL5 WT, C291Y, or C30W at baseline and following c-LTP (glycine) or c-LTD (NMDA) stimulation. The color bar indicates the intensity of CDKL5-PSD95 co-localization. For WT, a total of 25 dendrite segments from 5 neurons were analyzed in each group. A total of 15 dendrite segments from 3 neurons were analyzed for each group in C30W and C291Y plots. Two-tailed unpaired t-test or Mann-Whitney test with two tails was used. ***P < 0.001, ****P < 0.0001. Data are represented as mean ± SEM. **k**, Thermograph displaying the intensity of co-localization between Kalirin and PSD95 in CDKL5-deficient (493-stop) neurons replenished with CDKL5 WT, C291Y, or C30W at baseline and following c-LTP (glycine) or c-LTD (NMDA) stimulation. The color bar indicates the intensity of CDKL5-PSD95 co-localization. For each group, the same dendrite segments analyzed in (**j**) were used for analysis. Two-tailed unpaired t-test or two-tailed Mann-Whitney test was used. **P < 0.01, ***P < 0.001. Data are represented as mean ± SEM.

We next assessed the synaptic targeting of CDKL5^C30W^ and CDKL5^C291Y^, along with their impacts on the PSD localization of Kalirin7. When expressed in cultured CDKL5-deficient neurons, the CDKL5^C30W^ variant exhibited an enhanced enrichment in the PSD compared to CDKL5^WT^ (Fig. 7f, g). Neurons expressing CDKL5^C30W^ also exhibited increase in PSD size (Fig. 7h) and a higher Pearson’s correlation coefficient for Kalirin-PSD95 colocalization (Fig. 7i). These findings suggested that the p.C30W mutation results in the formation of more solid and condensed CDKL5-PSD95 co-condensates, which are associated with increased retention of Kalirin7 in the PSD due to reduced LLPS dynamics. In contrast, the CDKL5^C291Y^ mutant, which exhibits attenuated LLPS capacity, showed a significant decrease in synaptic localization (Fig. 7f, g) and a reduced Kalirin-PSD95 Pearson’s correlation coefficient (Fig. 7i), while the PSD area remained unchanged (Fig. 7h).

To determine whether these two mutations affect the activity-dependent synaptic translocation of CDKL5 and Kalirin7, we assessed the colocalization of CDKL5-Kalirin and Kalirin-PSD95 following c-LTP and c-LTD. CDKL5^WT^ exhibited increased CDKL5-PSD95 and Kalirin-PSD95 colocalization after c-LTP, while showing reduced CDKL5-PSD95 colocalization after c-LTD (Fig. 7j, k, left panel). In contrast, CDKL5^C30W^ completely abolished activity-dependent changes in CDKL5-PSD95 and Kalirin-PSD95 co-localization (Fig. 7j, k, middle panel), as evidenced by the colocalization coefficient relative to baseline (Ctrl groups). Furthermore, CDKL5^C291Y^ displayed a notable reduction in both CDKL5-PSD95 and Kalirin-PSD95 colocalization following c-LTP compared to CDKL5^WT^, likely attributable to its compromised LLPS capacity (Fig. 7j, k, right panel, Extended Data Fig. 6a, b).

Taken together, these results suggest that CDD-associated CDKL5 mutations alter the biophysical properties of LLPS, leading to dysregulated synaptic translocation of both CDKL5 and Kalirin7 in response to synaptic activity. The LLPS behavior of CDKL5 plays a crucial role in its synaptic localization, orchestrates synaptic structural plasticity, and regulates spine morphology (Fig. 8).

**Fig. 8.**
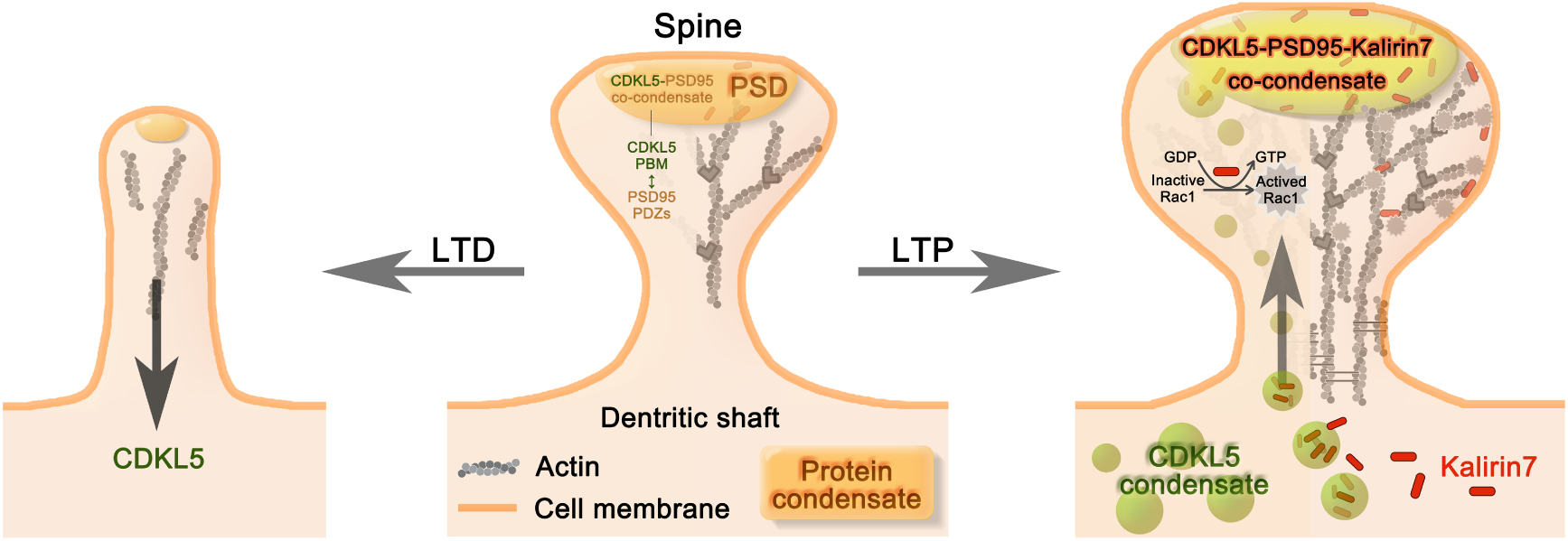
Model for functions of CDKL5 LLPS during synaptic plasticity. CDKL5 localizes within the PSD by forming co-condensates with PSD95 (middle). During LTP, CDKL5 disperses from PSD compartments, resulting in spine shrinkage (left); while during LTP, CDKL5 recruits and transports Kalirin7 into the PSD, activating Rac1 to assemble actin and promote spine enlargement (right).

## Discussion

The essential roles of CDKL5 in synaptic function are evidenced by the phenotypes of severe neuronal developmental delays and profound cognitive deficits in both CDKL5-deficient mice and patients bearing mutations in the *CDKL5* gene. Despite the strong correlation between CDKL5 deficiency and these phenotypic manifestations, the precise molecular mechanisms by which CDKL5 regulates synaptic activities remain largely unknown. In the current study, we have uncovered a novel mechanistic pathway by which CDKL5 governs synaptic plasticity. This mechanism is manifested as the bidirectional translocation of CDKL5 into and out of the PSDs in response to c-LTP and c-LTD, respectively. During c-LTP, CDKL5 specifically recruits and enriches Kalirin7 within the PSDs to modulate actin cytoskeleton reorganization, thereby facilitating activity-dependent dendritic spine enlargement. CDKL5 mutations linked to CDD disrupt the synaptic translocation of both CDKL5 and Kalirin7, leading to impaired synaptic plasticity, which may underlie the pathological mechanisms of CDD.

Since the identification of CDKL5 over two decades ago, it has been well established that its physiological functions in neural synapses are primarily mediated through kinase-dependent phosphorylation signaling cascades. For example, through chemical genetic approaches, microtubule-associated protein 1S (MAP1S) and EB2 have been identified as physiological substrates of CDKL5^38,39^. Importantly, phosphorylation of EB2 and MAP1S has been compromised in *Cdkl5* KO mice and neurons derived from CDD patients, along with abnormal microtubule dynamics and disrupted anterograde cargo trafficking in dendrites^38,40,41^. A recent study has identified the voltage-gated calcium channel Cav2.3 as a physiological substrate of CDKL5, and its loss of phosphorylation caused by deficiency in CDKL5 leads to increased neuronal excitability^42^. These findings highlight the importance of CDKL5 kinase activity in regulating synaptic development and functionality. However, the potential roles of CDKL5 that are independent of its enzymatic activity remain poorly understood. Moreover, little is known about how the C-terminus of CDKL5 contributes to its synaptic functions.

We discovered that CDKL5 possesses the intrinsic capacity to undergo LLPS both in vitro and in neurons. The IDR at the C-terminus, along with the KD of CDKL5, mediates this phase separation behavior. Importantly, we have established that the LLPS capability of CDKL5 is essential for the activity-dependent synaptic translocation of both CDKL5 and its interacting partner Kalirin7, a process critical for modulating synaptic structural plasticity. It is particularly noteworthy that while CDKL5 may execute its kinase function within dendritic spines, the kinase-dead mutation does not perturb its intrinsic LLPS capability. This observation strongly suggests that the LLPS-mediated synaptic targeting of CDKL5 and its partners is primarily governed by the phase separation properties of CDKL5 and operates independently of its catalytic activity. Recent investigations have revealed that the C-terminal domain of SynGAP, a synaptic GTPase-activating protein (GAP) that modulates Ras activity, plays a crucial role in mediating its LLPS with PSD95, rather than through its GAP activity. This domain is also essential for synaptic plasticity and cognitive behaviors^43^. We therefore propose a novel kinase-independent mechanism of synaptic regulation by CDKL5, which shares conceptual similarity with the phase separation-dependent functionality of SynGAP. Furthermore, the formation of CDKL5 enzymatic condensates may represent a novel regulatory mechanism for the spatial compartmentalization and concentration of limited CDKL5 molecules within specific subcellular domains, thereby facilitating rapid and spatially restricted catalytic activity. This phenomenon aligns with observations of other enzymatic condensates, including the synaptic GIT1/β-Pix complex, which has been demonstrated to undergo self-assembly independent of auxiliary scaffolding proteins ^44^. LLPS-mediated formation of enzyme condensates is likely to possess distinct characteristics with respect to enzyme kinetics, substrate accessibility, and catalytic efficiency. Future investigations are warranted to systematically elucidate how CDKL5 phase separation modulates kinase-dependent phosphorylation signaling cascades within synaptic compartments, particularly in relation to its role in synaptic plasticity and neuronal function.

Another important finding of this study is the identification of CDKL5-PSD95 co-condensate formation mediated through multivalent PDZ-PBM interactions. These heterotypic condensates are capable of recruiting Kalirin7, a well-characterized modulator of cytoskeletal dynamics, in an activity-dependent manner. Based on these findings and published studies, we propose a mechanistic model for synaptic plasticity regulation wherein LTP induction triggers CaMKII-mediated phosphorylation of SynGAP in an NMDA receptor-dependent manner, leading to its rapid dissociation from the PSD^17^. Concurrently, CDKL5 undergoes activity-dependent translocation into the PSD, forming the CDKL5-PSD95 co-condensates. This process is functionally complemented by recent findings demonstrating that transmembrane AMPAR regulatory proteins (TARPs), a class of AMPAR auxiliary subunits, also undergo phase separation with PSD95^45^. Consequently, the formation of CDKL5-PSD95 co-condensates is thus proposed to enhance the stabilization and clustering of AMPAR/TARP complexes within the PSD, thereby potentiating synaptic transmission. Furthermore, the LLPS-mediated recruitment of Kalirin7 by CDKL5 initiates Rac1 activation, which orchestrates actin cytoskeleton reorganization, ultimately contributing to dendritic spine enlargement and structural plasticity.

In summary, our findings establish a comprehensive molecular framework that delineates the role of CDKL5’s LLPS in synaptic function and its pathological implications when disrupted by CDD-associated mutations. This mechanistic understanding not only advances our fundamental knowledge of synaptic plasticity but also establishes a conceptual foundation for developing targeted therapeutic interventions specifically aimed at restoring normal LLPS and synaptic function of CDKL5 in patients with CDD.

## Materials and Methods

### DNA constructs

Human *CDKL5* DNA sequences, along with their truncated derivatives used in this study, were cloned from cDNA of 293T cells. The GFP-CDKL5 plasmid used in Fig. 1 and Extended Data Fig. 1 was cloned from cDNA of SD rat. The rat *PSD95* sequence used in HEK 293T cells was also cloned from SD rat cDNA. Rat *Kalirin7* sequences and their truncated derivatives were cloned from pEAK10-His-Myc-Kal7 (Addgene #25454). All CDKL5 mutations were generated using overlapping PCR, except for the PM mutation, which was synthesized by Sangon Biotech.

The C30W mutation was identified through an online survey of Chinese CDD families conducted by the authors. The reporting of the C30W mutation in this article was approved by the legal guardians of the related CDD patient and by the Ethics Committee of the Institute of Neuroscience, Center for Excellence in Brain Science and Intelligence Technology, Chinese Academy of Sciences (Research License CEBSIT-2024027).

Plasmids for transfecting 293T cells and primary neurons used the pEGFP-N1 vector as the backbone with the pCAG promoter. In the mEGFP-hCDKL5 constructs, A206 of EGFP was mutated to K to generate the mEGFP protein. For constructs containing red fluorescent proteins, EGFP was replaced with tdTomato, mRuby, or mCherry. For HA-Kalirin7, V5-PSD95, and other constructs with small tags, EGFP was replaced with the small tags. For lentivirus production, target sequences were inserted into a lentiviral packaging plasmid with pSyn as the promoter.

To express CDKL5^CTD^ protein in E. coli, the human *CDKL5* sequence was cloned into a vector containing an N-terminal TRX-His6 followed by an HRV 3C cutting site, using pET32M3C as the backbone. Various fragments of human PSD95 PDZ domains were amplified using standard PCR methods and also cloned into this pET32M3C vector. The full-length PSD95 protein was expressed using a pET32M3C-PSD95 FL plasmid constructed in a previous study^14^.

The generation of CDKL5 shRNA and scrambled shRNA constructs was described previously^12^. The target sequence of CDKL5 shRNA is GGAGCCTATGGAGTTGTAC, which has been extensively used ^12,36,46^.

### Protein expression and purification

Recombinant human PSD95 and CDKL5^CTD^ proteins were expressed in Escherichia coli Rosetta (DE3) cells. The PSD95 protein was expressed at 16°C for 18 hours, induced by isopropyl-β-D-thiogalactoside (IPTG) at a final concentration of 0.25 mM, and then purified by Ni2+-NTA agarose affinity chromatography and size-exclusion chromatography (with a column buffer containing 50 mM Tris pH 8.0, 100 mM NaCl, 1 mM EDTA, and 2 mM DTT). The CDKL5^CTD^ protein was expressed at 37°C for 3 hours and purified using TALON metal affinity resin (Clontech Laboratories, Inc. 635502), followed by size-exclusion chromatography (with a column buffer containing 20 mM HEPES pH 7.4, 450 mM NaCl, 1 mM EDTA, and 2 mM DTT). The TrX-His6 tag of purified proteins was cleaved by HRV 3C protease at 4°C overnight. Subsequently, a second round of size-exclusion chromatography was performed to remove the residual TrX-His6 tag and HRV 3C protease.

### Protein fluorescence labeling

The purified PSD95 proteins were transferred to a NaHCO_3_ buffer containing 100 mM NaHCO_3_ (pH 8.5), 100 mM NaCl, 1 mM EDTA, and 2 mM DTT using PD10 columns (Cytiva, 17085101). Subsequently, with a fluorophore to protein molar ratio of 1:1, Cy3 NHS ester (AAT Bioquest, no.141) was incubated with the corresponding protein at room temperature for 1 hour. Afterwards, the buffer of the labeled protein was changed back to 20 mM HEPES pH 7.4, 100 mM NaCl, 1 mM EDTA, and 2 mM DTT, while removing excessive dyes. BDP FL NHS ester (Lumiprobe, no. 11420) was used to label the CDKL5^CTD^ protein using the same procedure as the labeling of PSD proteins, except that the NaHCO_3_ buffer used in the labeling contained 450 mM NaCl. The fluorescence labeling efficiency of the labeled proteins was measured using a Nanodrop 2000 (Thermo Fisher), and the proteins were stored in aliquots at -80°C.

### In vitro phase transition and imaging assay

To form a CDKL5-PSD95 co-condensate, the two proteins were mixed at a stoichiometry of 1:10 or 1:5, with final concentrations ranging from 1-10 µM. The final concentration of each protein was indicated in the legend or figure. Typically, the ratio of fluorescence-labeled protein to unlabeled protein was 1:100. 20 mM HEPES buffer (pH 7.4) with higher NaCl concentration, and DTT were also added to the protein mixture to reach a final concentration of 20 mM HEPES, 150 mM NaCl, and 2 mM DTT.

A homemade chamber was created using a glass slide sandwiched between a coverslip and two layers of parafilm as a spacer for Nikon TiE imaging. The parafilm was cut into strips, placed parallel on the slide to form narrow channels that can indraft approximately 15-20 µl of protein mixture through capillarity. To secure the glass slide to the coverslip, the entire chamber was heated to 65°C on a metal bath for 10 seconds, which slightly melted the parafilm. Afterwards, the coverslip was gently pressed onto the glass slide to ensure adherence. Once the protein mixtures were injected, the chamber was placed on the microscope stage and allowed to settle for 10 minutes at room temperature with the coverslip facing down. All images were captured within 2 hours of mixing the proteins.

### Fluorescence polarization-based assay

The fluorescence polarization-based assay was conducted using a PerkinElmer LS-55 fluorimeter equipped with an automated polarizer at 25°C. In the assay, the commercially synthesized WT (NRSRMPNLNDLKETAL) and T958R (NRSRMPNLNDLKE**R**AL) mutant CDKL5-PBM peptides were labeled with fluorescein-5-isothiocyanate (FITC) (Invitrogen, Molecular Probe) at their N-terminal. The FITC-labeled WT or T958R CDKL5-PBM peptide was titrated with PSD95 PDZ proteins in a buffer containing 50 mM Tris (pH 8.0), 100 mM NaCl, 1 mM EDTA, and 2 mM DTT. The *K_d_* value was determined using a classical one-site specific binding model fitted with GraphPad Prism.

### 493-stop mice and HA-CDKL5 mice generating

The 493-stop mice (standard name C57BL/6-CDKL5^407+tm493stop.ZqxION^) were generated by combination of Cas9 with two sgRNA. sgRNA-1: TCTTTTGGACTGAGAAGGTG, and sgRNA-3: GTTTTGGACTTGGCTTCTTT, which deleted 2 base pairs at Lys410 position, led to premature terminal of CDKL5 protein at 493aa. HA-CDKL5 mice were generated by Shanghai Model Organisms Center, Inc. With homologous recombination induced through CRISPR/Cas9 technique, Kozak-3xHA sequence was inserted at start codon position of CDKL5 gene.

All experimental animals used in this study were housed in a 12-hour light/12-hour dark cycle with ad libitum access to food and water. All animal procedures were approved by the Institutional Animal Care and Use Committee of the Center for Excellence in Brain Science and Intelligent Technology of the Chinese Academy of Sciences.

### Cells culture and transfection

Cultures of hippocampal and cortical neurons were prepared from P0-P1 mice or rats, following previously described methods^47^. Neurons were transfected with plasmids using Lipofectamine 2000 (Invitrogen, no. 11668027) between DIV10 and DIV12. Lentivirus was transfected into neurons at DIV6 or 7 for a duration of 12 hours, followed by expression for at least 7 days. For live-cell imaging, neurons were planted on 35 mm glass-bottom dishes (FluoroDish FD35-100, World Precision Instruments, Inc.) coated with 0.1% poly-D-Lysine (Sigma-Aldrich, no. P0899). For biochemical tests, neurons were cultured in 6-well plates coated with 0.01% poly-D-Lysine. Neurons for immunocytochemistry assays were cultured on coverslips coated with 0.1% poly-D-Lysine. HEK 293T cells (from ATCC) were cultured in DMEM media containing 10% fetal bovine serum in a 37°C incubator with 5% CO_2_. Cells for biochemical tests were cultured in 6-well or 12-well plates, while cells for immunocytochemistry assays were cultured on coverslips. Plasmids were transfected into 293T cells using Lipo2000 and then expressed for 24 hours.

### Immunocytochemistry

For the immunocytochemistry assay, cells were washed with PBS and fixed in cold methanol for 15 minutes. After fixation, the cells were washed once with PBS, then blocked and permeabilized in PBS containing 3% BSA and 0.5% Triton X-100 for 30 minutes at room temperature. They were then incubated with the primary antibody overnight at 4°C. The following day, cells were washed with PBS three times and incubated with Alexa Fluor-conjugated secondary antibodies for 1 hour at room temperature. After washing off the secondary antibodies once, cell nuclei were stained with Hoechst at 0.5 mg/ml in PBS for 10 minutes at room temperature, if nuclear staining was required. After an additional wash, the cells were mounted in antifade resin glue.

Generally, the primary antibodies were diluted at ratios ranging from 1:200 to 1:500 for this assay, while the secondary antibodies were diluted at a 1:200 ratio. The primary antibodies used included chicken anti-GFP (Invitrogen, no. A10262), rat anti-CDKL5 (Abmart Inc, no. X-E2E1S0), rabbit anti-Piccolo (Proteintech, no. 18156-1-AP), mouse anti-PSD95 (Abcam, no. ab192757), rabbit anti-Kalirin (Sigma-Aldrich, no. 07-122, targeting residues 517-976 of rat Kalirin and recognizing multiple isoforms), mouse anti-Rac1 (Proteintech, no. 66122-1-Ig), rabbit anti-V5 (CST, no. 13202), and rat anti-HA (Roche, no. 11867423001). For secondary antibodies, Invitrogen Alexa Fluor products were used, including 405-tagged (no. A48258), 488-tagged (no. A32790), and 555-tagged (no. A32794) anti-rabbit IgG; 488-tagged (no. A21208) and 546-tagged (no. A11081) anti-rat IgG; 405-tagged (no. A31553), 555-tagged (no. A32773), and 647-tagged (no. A21236) anti-mouse IgG; and 488-tagged (no. A32931) anti-chicken IgG.

### Fluorescence recovery after photo-bleaching assay (FRAP assay)

293T cells were cultured in 35 mm glass-bottom dishes and transfected as previously described. The culture medium was replaced with a transparent, prewarmed artificial cerebrospinal fluid (ACSF: 125 mM NaCl, 2.5 mM KCl, 25 mM HEPES pH 7.3, 33 mM D-glucose, 2 mM CaCl_2_, 1 mM MgCl_2_), and the image acquisition was carried out in a 37 °C chamber with 5% CO_2_. Time-lapse images were captured at 2-second intervals for 1 minute and then aligned using the ImageJ plugin MultistackReg. Five frames were acquired as a baseline before the start of photobleaching, and the average fluorescence intensities of puncta were normalized to 100%. The GFP signal was bleached using a 488-nm laser beam. The outlines of puncta were manually drawn, and puncta that moved out of the focal plane or merged with others were excluded from the data analysis.

### Chemical LTD and chemical LTP

The protocols of NMDA-induce chemical LTD and 0 Mg^2+^/glycine -induced chemical LTP were modified from a recent study^17^.

NMDA-induced chemical LTD: for biochemistry experiments, neurons were incubated in cultured medium with 50 µM NMDA for 10 min before harvested; for imaging, neurons were incubated in artificial cerebrospinal fluid (ACSF: 125 mM NaCl, 2.5 mM KCl, 25 mM HEPES pH 7.3, 33 mM D-glucose, 2 mM CaCl_2_, 1 mM MgCl_2_, 1 µM TTX) for 20 min prior to switch to LTD induction solution (ACSF with 0 mM MgCl_2_, 20 µM NMDA). After incubation in induction solution for 10 min, the solution was switched back to ACSF for indicated time.

0 Mg^2+^/glycine-induced chemical LTP: neurons were incubated in ACSF (125 mM NaCl, 2.5 mM KCl, 25 mM HEPES pH 7.3, 33 mM D-glucose, 2 mM CaCl_2_, 1 mM MgCl_2_, 0.5 µM TTX, 1 µM Strychnine, 20 µM Bicuculline) for 20 min, then applied with LTP induction solution (ACSF with 0 mM MgCl_2_, 100 µM glycine) for 5 min (mouse neurons) or 10 min (rat neurons). Solution was switched back to ACSF for wash out at indicated time.

### Imaging and data analysis

Images of fixed cells were acquired using a Nikon TiE or NiE laser scanning microscope with either a 60x/1.4 NA oil immersion objective lens, or a 40x/0.95 NA air objective lens. Live-cell imaging, including the FRAP assay, was conducted using a Nikon A1R confocal microscope with a 60x/1.4 NA oil immersion objective lens. Generally, for 293T cell samples, 60x z-stack images were collected at a resolution of 1024x1024 pixels, averaged twice, and taken at 0.3 µm intervals; 40x z-stack images were collected at a resolution of 1024x1024 pixels, and taken at 1 µm intervals, without averaging. For primary neuron samples, z-stack images were collected at the same resolution, averaged twice, and taken at 0.2 µm intervals. For in vitro LLPS imaging, z-stack images were collected at a resolution of 1024x1024 pixels and taken at 1 µm or 0.5 µm intervals without averaging. The results are presented as z-stacked images unless a specific image is identified as a single slice in the figure legends. Overexposure was avoided during image acquisition whenever possible.

Images were analyzed using ImageJ. The cluster intensity of PSD95 and CDKL5 in Fig. 3d was measured in a ROI defined by a manually set threshold. Pearson’s coefficient, calculated using the colocalization threshold ImageJ plug-in, was employed to analyze the co-localization of two proteins in mice primary neuron samples. The partition coefficient (Fig. 3d) was used to analyze the preferred PDZ domain of PSD95 binding to the PBM of CDKL5 by measuring the concentration ratio between the macromolecule inside the dense phase and outside the light phase of the biomolecular condensate. A threshold was set to define regions of CDKL5 puncta across all groups, excluding the cell nuclear region. The mean grey value of PSD95 within the CDKL5 puncta regions represented the dense phase concentration (C dense), while the mean grey value of cytoplasmic PSD95 outside the CDKL5 puncta regions was defined as the light phase concentration (C light). The partition coefficient w then calculated using the following formula:

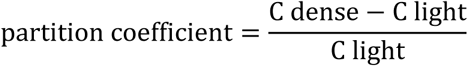

### Brain tissue lysate preparation

Mice tissues were lysed in RIPA buffer (50 mM Tris-HCl, pH 7.4, 150 mM NaCl, 1% TritonX-100, 0.5% Sodium deoxycholate, 0.1% SDS, 1 mM EDTA) with 1 x protease inhibitors (Acmec, no. AC13365) and phosphatase inhibitors (1 mM NaF and 1mM Na_3_VO_4_) and homogenized with a frozen ball mill (Shanghai Jingxin industrial development company). The homogenates were lysed for 20 min on ice then centrifuged at 14,000 x g for 20 min at 4°C. The protein concentration of the supernatants was then determined using a BCA protein assay kit (Tiangen, no. PA115).

### CDKL5^CTD^ condensate pull-down assay

Purified CDKL5^CTD^ protein was mixed with PEG 8000 to achieve a final concentration of 3% to induce LLPS. The mixture was allowed to settle gravitationally in a 96-well plate for 30 minutes at room temperature, resulting in droplets fusing into a protein liquid layer at the bottom. After discarding the CTD dilute phase, WT mouse brain lysate was incubated with the CTD protein layer for 30 minutes at room temperature. A portion of the supernatant was reserved as sample “S” Following a gentle rinse with PBS, the CTD layer was resuspended with RIPA strong lysis buffer (Beyotime, no. P0013B) as sample “P”. This sample was then subjected to gel electrophoresis and silver staining. Protein bands that exhibited significant enrichment in sample “P” compared to samples “S” (approximately at the 250 kDa position and higher), along with the corresponding gel segment from the brain lysate lane, were excised from the gel and analyzed using mass spectrometry. Based on the mass spectrometric analysis (by Shanghai Applied Protein Technology Co.), proteins that exhibited a higher concentration in sample “P” than in the brain lysate were considered potential targets of the CDKL5 condensate.

### Gel electrophoresis and Western blotting

Protein samples were separated using a 9%–12% SDS-polyacrylamide gel, with the gel concentration chosen based on the molecular weight of the target proteins. For Western blotting, the samples were transferred onto a PVDF membrane (Millipore, no. IPVH00010) following gel electrophoresis. After blocking with 5% non-fat milk in Tris-buffered saline with 0.1% Tween^®^ 20 detergent (TBST) at room temperature for 1 hour, the membranes were incubated overnight at 4°C with primary antibodies, followed by a 2-hour incubation at room temperature with corresponding HRP-conjugated secondary antibodies. For CDKL5 blotting, sheep anti-CDKL5 (Protein Phosphorylation and Ubiquitylation, no. 3065179) was used. In Extended Data Fig. 3a, rabbit anti-PSD95 was used (CST, no. 3450). Bands were visualized using enhanced chemiluminescence (ECL) and quantified with ImageJ software.

### Structure prediction

The structures of the PDZ-PBM complexes were predicted utilizing the AlphaFold3 server (https://alphafoldserver.com/). The amino acid sequences for human PSD95 (UniProt ID: P78352) and CDKL5 (UniProt ID: O76039) were obtained from the UniProt database. All predicted template modeling (pTM) scores for the complex structures exceeded 0.7, with interface pTM (ipTM) scores of 0.75 (PDZ1-PBM), 0.77 (PDZ2-PBM), and 0.82 (PDZ3-PBM). These scores indicate high-confidence model predictions according to established AlphaFold3 evaluation metrics. Structural visualization and analysis were conducted using PyMOL.

### Statistical analysis

Values are reported as mean ± SEM, unless otherwise specified. The normal distribution of values was verified using the Shapiro-Wilk test. To determine the significance of differences between two groups, two-tailed Student t-test was conducted. If the data did not follow a normal distribution, two-tailed Mann-Whitney U test was used. Kolmogorov-Smirnov test was used to measure significance of differences between two cumulative curves. The statistical significance was defined as *p < 0.05, **p < 0.01, ***p < 0.001, ****p < 0.0001.

## Acknowledgments

We thank Yi-Dian Zhang for support in constructing the HA-CDKL5 mice; Dr. Steven Boeynaems for advice in PM mutation construction; Dr. Qian Hu, Dan Xiang, and Xu-Xin Chen from the Center for Excellence in Brain Science and Intelligent Technology of the Chinese Academy of Sciences Optical Imaging Core Facility for assistance with microscopy and image data analysis; Ling Han from the Animal Facility for help with animal care; Yi-Wen Zhang, Fan Yang, and Tang Wei from the Gene Editing Core Facility for lentivirus production; Song-Lin Qian from the Molecular and Cellular Biology Core Facility for assistance with preliminary experiments in protein expression and purification; and the members of the Z.-Q.X. laboratory for their helpful discussions.

This work was supported by grants from Innovation of Science and Technology 2030-Major Project “platform of nonhuman primate models” (Grant No.2021ZD0200900 to Z.-Q.X.), National Natural Science Foundation of China (82021001 to Z.-Q.X.), Shanghai Municipal Science and Technology Major Project to Z.-Q.X.; STI2030-Major Project (2022ZD0214400) to J.-W. Z., National Natural Science Foundation of China (32470728) to J.-W. Z., Natural Science Foundation of Chongqing, China (CSTB2023NSCQ-JQX0031) to J.-W. Z.; STI2030-Major Project (2022ZD0214200), and National Natural Science Foundation of China (32371075) to Y.-C.Z..

## Author contributions

M.-J.L., M.-J.Z., and Z.-Q.X. conceived this study. M.-J.L. and J.-W. Z. designed the in vitro LLPS experiments with the guidance of M.-J.Z., M.-J.L. and Y.-C.Z. designed the experiments involving cultured neurons in Fig. 2-Fig. 7, with M.-J.L. performed these experiments. D.L. and Z.-A.Z. discovered the activity-dependent trafficking of CDKL5 and contributed data related to Fig. 1. M.-J.L. and Z.-A.Z. conducted experiments using 239T cells together. M.-J.Z. provided plasmids and purification protocols for PSD proteins. J.-C. W. provided all proteins used in the in vitro LLPS assays, conducted the fluorescence polarization-based assay, and generated protein complex structural models. M.-J.L. and Z.-A.Z. analyzed the images. M.-J.L., J.-W. Z., and Y.-C.Z. wrote the manuscript with input from all authors. Z.-A.Z. and Z.-Q.X. revised the manuscript. Y.-C.Z., J.-W. Z., and Z.-Q.X. supervised the study.

---

**Extended Data Fig. 1.**
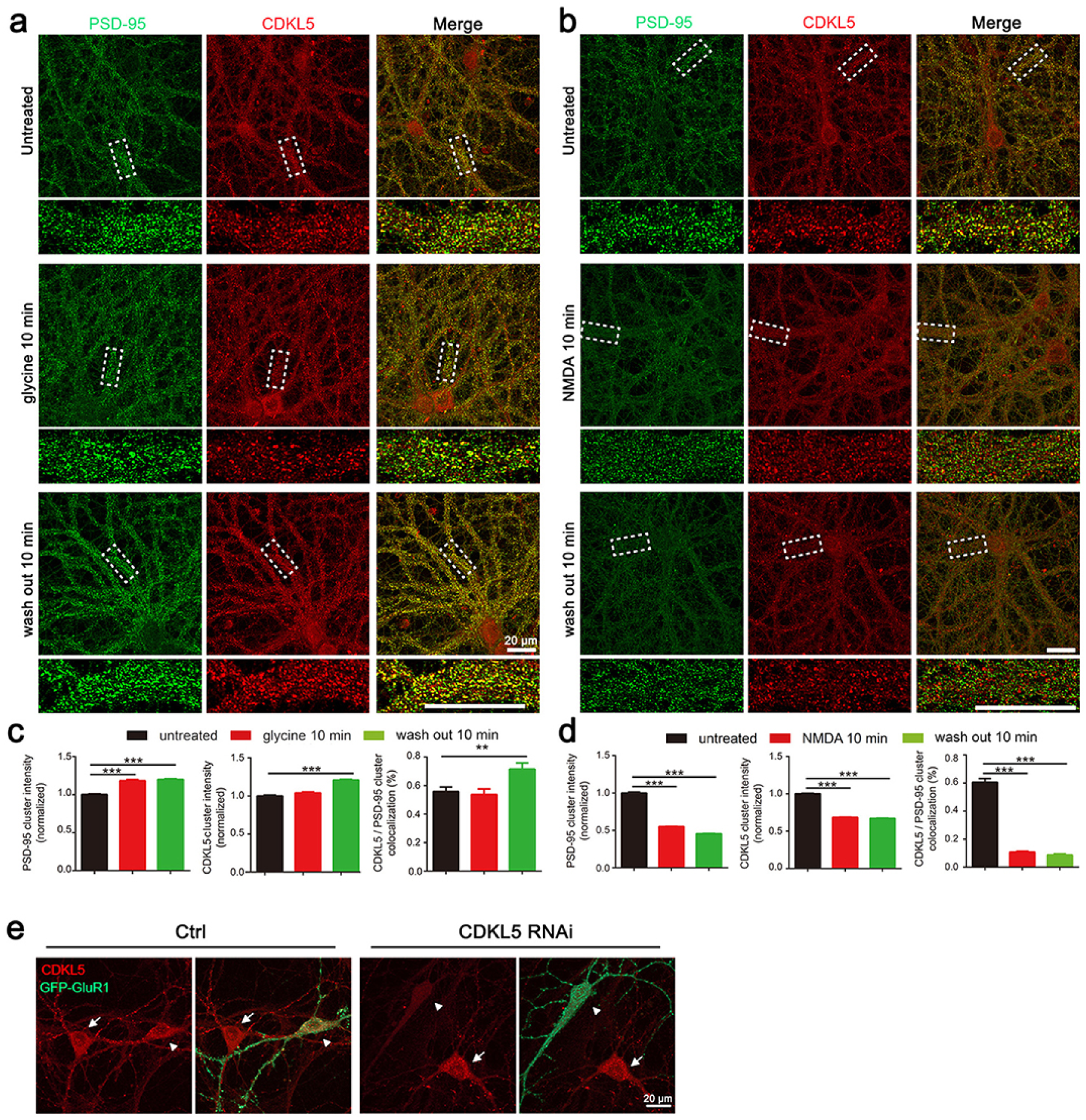
c-LTP stimulation increases the co-localization of CDKL5 with PSD95, while c-LTD treatment reduces it. **a**, **b**, Representative ICC images showing the endogenous distribution of CDKL5 and PSD95 before and after glycine (**a**) or NMDA (**b**) stimulation. Dashed boxes indicate the zoomed-in regions. Scale bar: 20 μm. **c**, **d**, Quantification of CDKL5 and PSD95 signal intensity, and percentage of CDKL5-PSD95 colocalization during c-LTP (**c**) and c-LTD (**d**). **e**, Representative images demonstrating efficient CDKL5 knockdown by shRNA in transfected neurons. White triangles indicate shRNA-transfected neurons marked with GFP-GluR1 (green), while the white arrows indicate untransfected neurons. In the control group (left panel), scrambled shRNA was delivered, showing no knockdown effect, whereas CDKL5 shRNA effectively reduced CDKL5 protein levels (right panel).

**Extended Data Fig. 2.**
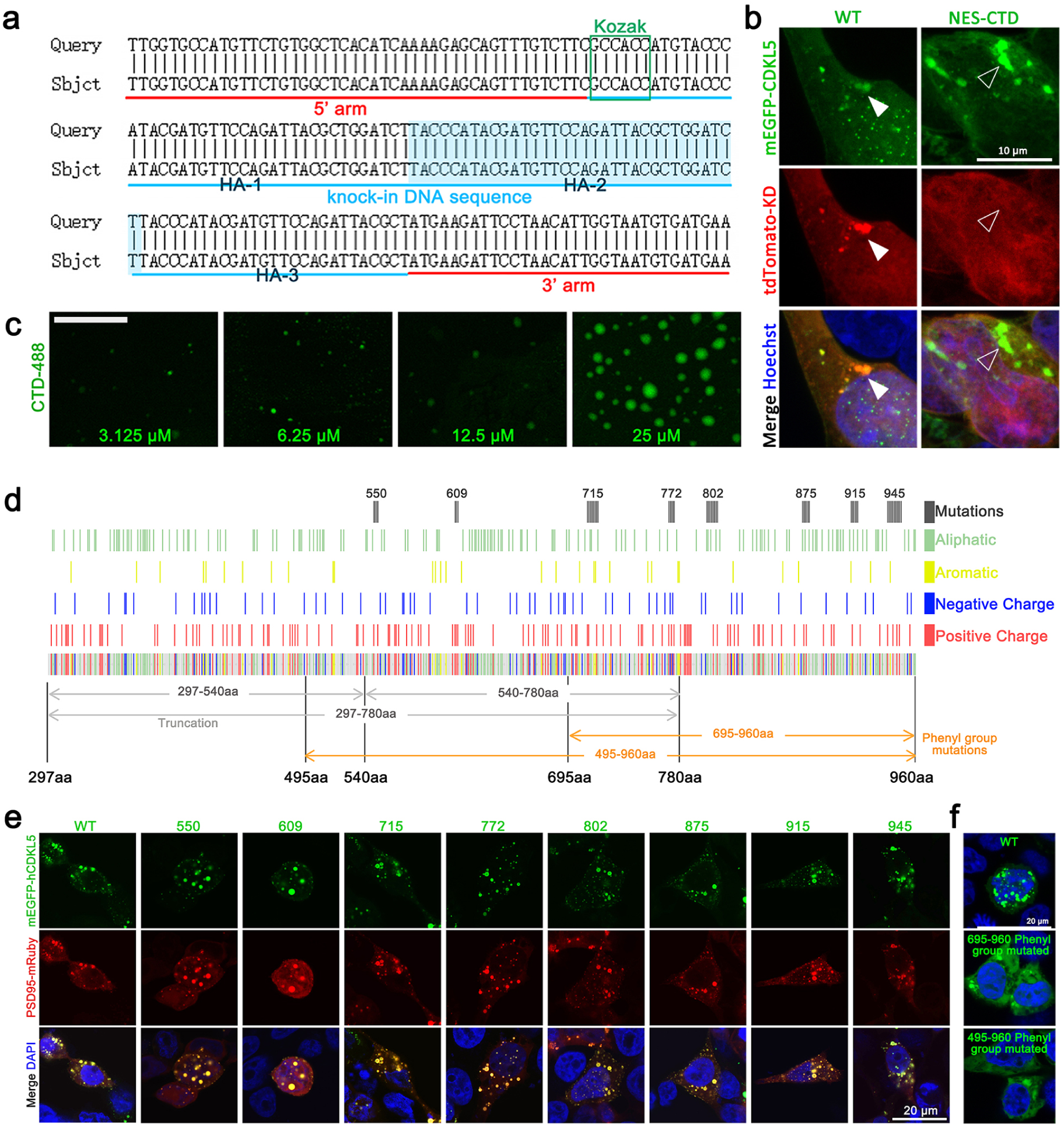
Mutating aromatic amino acids of the CTD reduces the LLPS of CDKL5, while mutating clustered polar residues fails to inhibit CDKL5’s LLPS. **a**, Strategy for generating HA tag knock-in mice (HA-CDKL5 mice). The query represents the designed sequence, while the subject sequence shows the actual sequencing results of the HA-CDKL5 mice. **b**, Images showing tdtomato-CDKL5-KD can be enriched by mEGFP-CDKL5-WT but not by NES-mEGFP-CDKL5^CTD^. The hollow and solid arrowheads indicate NES-CTD and WT puncta, respectively. **c,** Representative images of protein droplets formed by the CTD protein, with concentration gradients of 3.125 μM, 6.25 μM, 12.5 μM, and 25 μM. Scale bar: 20 μm **d**, Diagram illustrating the distribution of different groups of residues along the CTD of CDKL5. The indicated truncated mutations shown in this figure were used in subsequent experiments. **e**, Co-expression of PSD95-mRuby with mEGFP-tagged WT or mutants of a closely clustered group of hydrophobic, charged, or aromatic residues at various positions, including 550 (R^547^NNR to ANNA), 609 (H^609^RH to AAA), 715 (R^710^RVGSSFYRV to AAAGSAAAA), 772 (K^772^EKEK to AEAEA), 802 (L^801^LTLQKSIH to AATAQASAA), 875 (I^874^RIHPL to AAAAPA), 915 (W^911^HVSSV to AAASSA), 945 (H^939^PYNRTNRSSRM to APANATNASAA). Scale bar: 20 μm. **f**, Representative images showing LLPS of mEGFP-WT or mEGFP-tagged CDKL5 with phenyl group mutations at positions 695-960 aa or 495-960 aa. Scale bar: 20 μm.

**Extended Data Fig. 3.**
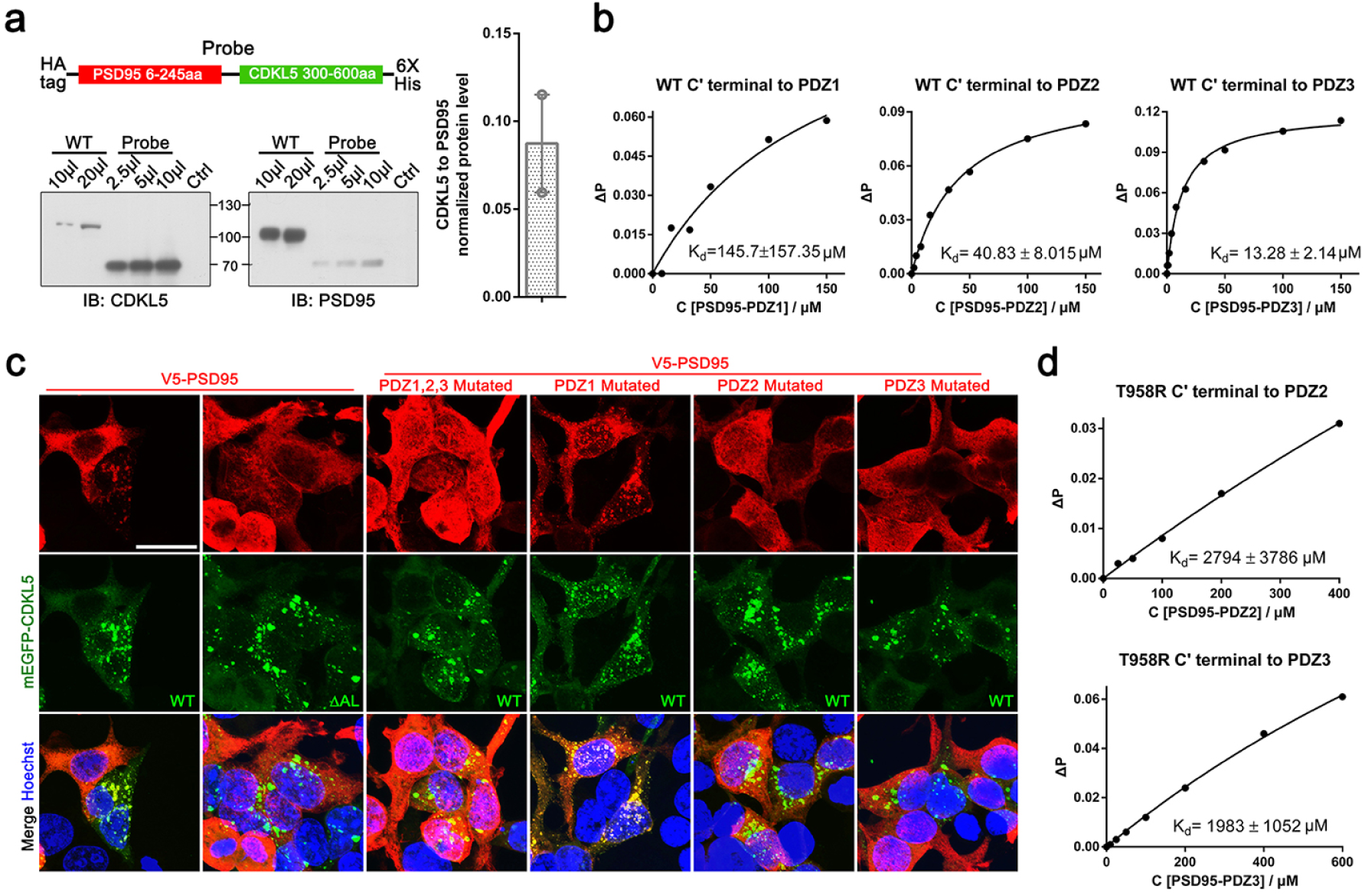
PBM of CDKL5 interacts with multiple PDZ domains of PSD95. **a**, Diagram of chimeric protein “Probe” construct and immunoblots from WT mouse brain lysate. The “Probe” contains binding regions for both the PSD95 and CDKL5 antibodies. The ratio of endogenous CDKL5 to PSD95 was calculated by normalizing the intensity of CDKL5 or PSD95 bands to the “Probe” band, yielding a ratio of approximately 1:10. Two replicates were analyzed in the plot at right panel. **b**, Plots of fluorescence polarization-based assays measurements calculating the *K_d_* between CDKL5 PBM and each PDZ domain of PSD95. **c**, Representative images of 293T cells co-expressing mEGFP-CDKL5 WT and V5-PSD95 PDZ domain mutations, corresponding to the PSD95 partition coefficient calculated in Fig. (**3d**). Scale bar: 20 μm. **d**, Plots of fluorescence polarization-based assays measurements calculating the *K_d_*between T958R and the PDZ2 or PDZ3 domains of PSD95.

**Extended Data Fig. 4.**
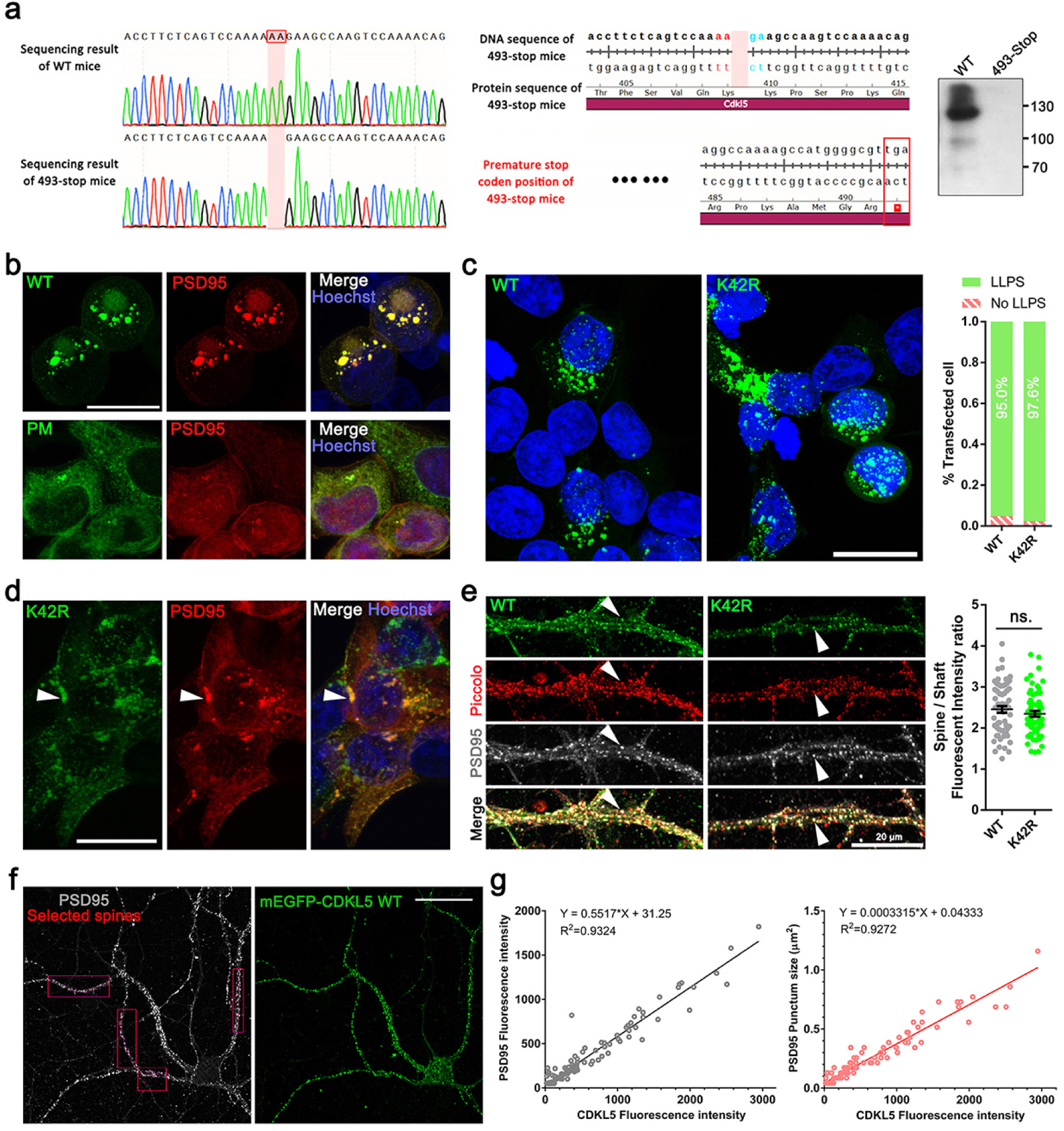
K42R shows minimal impact on CDKL5 LLPS, and CDKL5 intensity is correlated with PSD size and density. **a**, Diagram illustrating the DNA genome (left panel) and protein sequences (middle panel) at the mutated position in the 493-stop mice, and immunoblot of CDKL5 protein using brain lysates from WT and 493-stop mice (right panel). The standard name for this mouse line is C57BL/6-CDKL5^407+tm493stop.ZqxION^, abbreviated as 493-stop. **b**, Representative images showing mRuby-PSD95 puncta enriched in CDKL5-WT, but not in CDKL5-PM mutation. Scale bar: 20 μm. **c**, Images and quantification of the percentage of LLPS 293T cells expressing mEGFP-CDKL5 WT or K42R (using the same WT data as in Fig. (**2e, g**) to calculate the percentage of LLPS cells). Scale bar: 20 μm. **d.** Images of 293T cells co-expressing V5-PSD95 and mEGFP-CDKL5-K42R showing partial colocalization. White pyramids indicate CDKL5-PSD95 colocalized puncta. Scale bar: 20 μm. **e,** Images of 493-stop neurons expressing CDKL5 WT or K42R, with white pyramids indicating PSD-localized CDKL5. The synaptic location of CDKL5 was measured by the spine-shaft intensity ratio. Piccolo and PSD95 were used to define the spine region. For each group, 60 spines were analyzed, and unpaired two-tailed t-test was used for the analysis. Data are represented as mean ± SEM. **f**, Images of 493-stop neurons expressing mEGFP-CDKL5 WT were transfected with lentivirus at DIV 7 and fixed for ICC at DIV 14. The red boxes indicate the regions of interest (ROIs) where spines were selected for analysis performed in (**g**). Scale bar: 50 μm. **g**, Results of linear fitting for fluorescence intensity of CDKL5 versus colocalized PSD95 intensity (left), and puncta size (right). Each dot represents an individual CDKL5-PSD95 punctum, with a total of 100 spines included.

**Extended Data Fig. 5.**
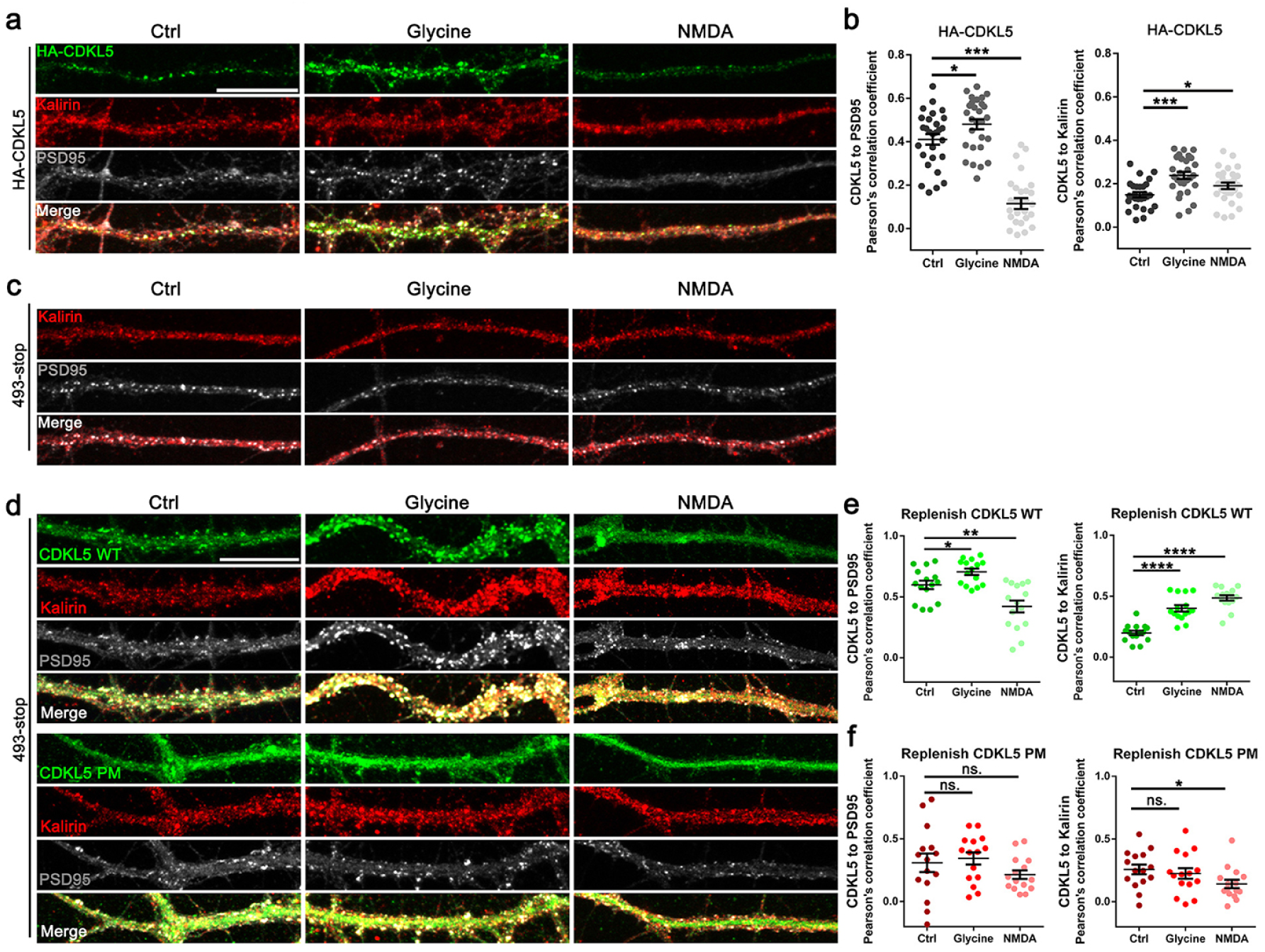
Co-localization of CDKL5-PSD95 and CDKL5-Kalirin are both response to neuronal activities. **a,** Corresponding fluorescence images of HA-CDKL5 neurons for thermography shown in Fig. (**6e**), scale bar: 20 μm. **b,** Pearson’s correlation coefficient of CDKL5-PSD95 and CDKL5-Kalirin in HA-CDKL5 neurons. A total of 15 dendrite segments from 3 neurons for each group were analyzed. Unpaired t-test with two tails was used. *P < 0.05, **P<0.01, ****P < 0.0001. Data are represented as mean ± SEM. **c,** Corresponding fluorescence images of 493-stop neurons for thermography shown in Fig. (**6f**), sharing the same scale bar with (**a**). **d**, Fluorescence images of 493-stop neurons correspond to the thermography shown in Fig. (**6g, h**), Scale bar: 20 μm. **e, f,** The Pearson’s correlation coefficient of CDKL5-PSD95 and CDKL5-Kalirin in 493-stop neurons replenished with CDKL5 WT or PM showed in Fig. (**6g, h**). A total of 15 dendrite segments from 3 neurons for each group were analyzed. Unpaired t-test with two tails was used. *P < 0.05, **P < 0.01, ****P < 0.0001. Data are represented as mean ± SEM.

**Extended Data Fig. 6.**
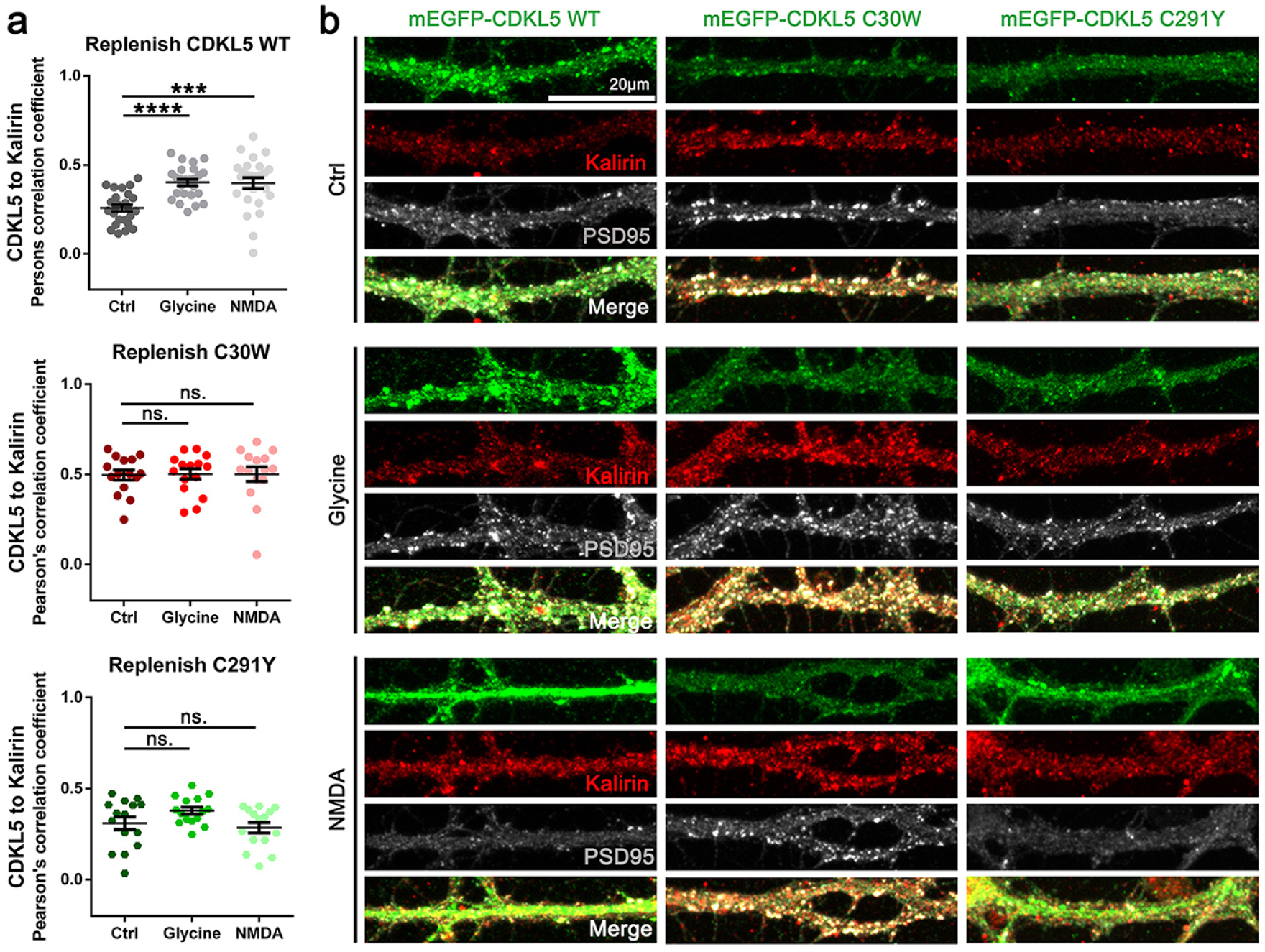
CDD-associated point mutations impair activity-dependent translocation of CDKL5 and co-localization with Kalirin. **a**, Pearson’s correlation coefficient of CDKL5 to Kalirin in CDKL5-deficient (493-stop) neurons replenished with CDKL5 WT, C30W, or C291Y. For each group, the same dendrite segments analyzed in Fig. 7 (**j, k**) were used for analysis. Two-tailed unpaired t-test or two-tailed Mann-Whitney test was used. ***P < 0.001, ****P < 0.0001. Data are represented as mean ± SEM. **b**, Corresponding fluorescence images of thermographs shown in Fig. 7 (**j, k**).

